# Environmentally mediated interactions predict community assembly and invasion success in a gut microbiota SynCom

**DOI:** 10.1101/2024.07.15.603569

**Authors:** P.T. van Leeuwen, P. Gadaleta, S. Brul, J. Seppen, M.T. Wortel

**Affiliations:** Molecular Biology and Microbial Food Safety, Swammerdam Institute for Life Sciences, University of Amsterdam, Amsterdam, The Netherlands; Tytgat Institute for Liver and Intestinal Research, Amsterdam University Medical Centers, Location AMC, 1105 BK Amsterdam, The Netherlands

## Abstract

The gut microbiome plays a crucial role in host homeostasis, with implications for nutrition, immune development, metabolism, and protection against pathogens. Disturbance of the microbiome by microbial invasion can be negative or positive: invasions of opportunistic pathogens can cause disease while dysbiotic states need invasions to recover. However, the complexity of the microbiome challenges our understanding of what factors determine the ability of microbes to invade. In this study we measure interactions between members of a SynCom of prominent gut bacteria with supernatant assays. We measure relative abundances of growth of co-cultures up to four species to validate a generalised Lotka-Volterra model parameterized with these supernatant assays. We predict differential invasion profiles of the opportunistic pathogens *Escherichia coli* and *Bacteroides ovatus* based on their monoculture growth profiles and interactions with other species. With experimental data of both predicted invasion and resistant communities we confirm model predictions of invasion success. Our model shows that both negative interactions within a community and neutral to positive interactions with the invading species promote invasion success, but the interactions towards the invading species dominate. Our validated approach opens the way for testing of interactions of human gut microbiome species, thereby developing inventions to avoid pathogenic overgrowth and therapies to enhance health-benefitting invasions.

**Importance:** The stability of the human gut microbiome is crucial for host health, with opportunistic pathogen invasions causing diseases and healthy strain replacements needed for recovery. The microbiota’s complexity complicates understanding its stabilizing mechanisms. This study uses a 10-species synthetic community of common gut microbiota to predict stable communities and invasion success. We employ supernatant experiments to parameterize a computational model, accurately predicting community composition and invasion success of Escherichia coli and Bacteroides ovatus. Our findings show that interactions within the resident community and with the invader are important, but the latter dominate. These results pave the way for larger-scale studies to characterize gut microbiome interactions and properties that resist invasions, potentially benefiting health through improved probiotics and faecal microbiota transplants.

## Introduction

The human gut microbiota is important in host health, including resistance to invasion by opportunistic pathogens. A healthy microbiota is often characterised by high species diversity and temporal stability, while an unstable or disrupted microbiota is linked to a range of health issues, including inflammatory bowel disease and other gastrointestinal disorders (Ley et al. 2005; 2005; Ridaura et al. 2013; Carding et al. 2015; Chang and Lin 2016; Saffouri et al. 2019; Fassarella et al. 2021; Sultan et al. 2021; Santana et al. 2022).

Overgrowth of opportunistic pathogens can lead to diseases such as *Clostridioides difficile* infection, causing antibiotic-associated diarrhea and pseudomembranous colitis (Mullish and Williams 2018); Candida overgrowth, causing gastrointestinal candidiasis and potentially invasive candidiasis (Alonso-Monge et al. 2021); small intestinal bacterial overgrowth, causing bloating, malabsorption and nutrient deficiencies (Ghoshal et al. 2017); expansion of Proteobacteria/Enterobacteriaceae, driving intestinal inflammation and increasing the risk of bacterial translocation, sepsis or worsened inflammatory bowel disease (Lupp et al. 2007); and Fusobacterium nucleatum overgrowth, which has been linked to colorectal cancer progression (Shang and Liu 2018). Strategies to treat some of these diseases by faecal microbiota transplantation or targeted probiotic administration rely on engraftment, the successful integration of beneficial microbes into existing communities, which might be aided by previous administration of antibiotics (Shogbesan et al. 2018). Since the efficacy of intervention such as FMT in inflammatory bowel disease is limited (Arora et al. 2023), it is crucial to understand how microbial species can successfully engraft in existing gut microbial communities. Invasions in microbial communities have been studied with both theoretical approaches and experimental systems, aiming to understand which community properties affect invasion.

Invasion can be classified as successful, with the resident community retaining its diversity (augmentation) or not (displacement), or as unsuccessful, with maintenance of the resident community (resistance) or not (disruption). In theoretical ecology, larger community size and stronger competition have been shown to resist invasion (Case 1990). More recently, ecological concepts are applied to microbiology and human microbiota (Costello et al. 2012; Bakkeren et al. 2025). For example, increased diversity has been linked to invasion resistance in microbial communities (Mallon *et al*. 2015). This relationship has mixed experimental support, leading to the prediction by Hu *et al*. (2025) that it is the dynamic regime of the resident community—oscillations versus stable states—that can predict invasion success. The outcome of microbial invasion is not solely determined by antagonistic interactions; facilitative interactions, where resident microbes assist invaders by altering environmental conditions or sharing metabolites, can promote successful invasions with either augmentation or displacement (Zhu and Momeni 2024). Together, these studies highlight how important the specific ecological and interactional context is for assessing invasions. Here, we investigate which properties affect invasion in a community of common human gut microbes using a combined experimental and computational approach.

Synthetic communities (Syncoms) are useful tools to understand complex communities, capturing some of the complex interactions but retaining enough simplicity for experimental tractability (Shetty et al. 2022; van Leeuwen et al. 2023). Syncoms of the human gut microbiota are used in many studies (Petrof et al. 2013; Desai et al. 2016; Mark Welch et al. 2017a; D’hoe et al. 2018; Venturelli et al. 2018; Ziesack et al. 2018; El Hage 2019; Gutiérrez and Garrido 2019). We selected 10 abundant human gut microbes to cover a broad range of functions of the microbiome. These species, *Akkermansia muciniphila, Bacteroides ovatus, Bacteroides thetaiotaomicron, Bifidobacterium adolescentis, Blautia obeum, Lachnospiraceae spp., Lactobacillus johnsonii, Faecalibacterium prausnitzii, Roseburia intestinalis* and *Escherichia coli*, spread over the phyla present in the western human gut and have been characterized in terms of their association with health or disease (Rinninella et al. 2019; Pittayanon et al. 2020; Haneishi et al. 2023) (Supplementary Figure 1). They exhibit complementary metabolic capacities centered on the fermentation of polysaccharides and mucins, leading to the production of short-chain fatty acids (SCFAs) such as acetate, propionate, and butyrate (Flint et al. 2012; Louis and Flint 2017) and contain obligatory anaerobes (members of the *Bacteroides, Blautia, Lachnospiraceae, Faecalibacterium*, and *Bifidobacterium*) as well as facultative anaerobes capable of growth under microaerophilic condition (*Lactobacillus johnsonii* and *Escherichia coli*) (Donaldson et al. 2016)). Altered abundance or loss of these species has been linked to metabolic and inflammatory disorders including obesity, type 2 diabetes, and inflammatory bowel disease (Qin et al. 2010; Derrien et al. 2017).

Interactions between microbes can originate from mainly metabolic interactions, toxin production and physical interactions. Supernatant experiments, where environmentally mediated interactions are considered, have shown to capture an important part of species interactions (de Vos et al. 2017; Vetsigian et al. 2011; Goldman et al. 2025; Dos Santos et al. 2022). This agrees with the observation that species interactions can for a large part be described by a kinetic model when metabolite measurements are included (D’hoe et al. 2018). With only species measurements, species interaction models such as Lotka-Volterra competition models or its generalizations can be used to predict species dynamics. These models come with inherent limitations, such as limited predictability of batch cultures and the exclusion of higher level interactions (Picot et al. 2023; Momeni et al. 2017). Despite these shortcomings, they have been used to predict dynamics in a human gut microbiota community and cocultures of pathogens (Venturelli et al. 2018; Ye et al. 2014).

Here, we use conditioned media assays, a form of supernatant assays, to describe the environmentally mediated species interaction structure of our SynCom. Next, we use a generalized Lotka-Volterra model to predict co-culture growth and validate these results with co-cultures of up to four species. Subsequently, we predict and validate species invasions in stable communities and elucidate the most important properties of the invader and resident community that predict invasion success.

## Results

### Negative interactions dominate between SynCom members

We generated growth curves in a conditioned medium composed of 40% spent media of the donor species bacterial species and 60% fresh YCFA to assess species growth characteristics and interactions. Overall, we see that negative interactions, which could be caused by e.g. resource competition or toxin production, dominate between the tested species (Figure 1 and Supplementary Figure 2AB). Although the effects on growth rate and final population density correlate (Supplementary Figure 3), the effects of conditioned media on growth rate are less negative and more often positive than on maximum population density, suggesting that there are growth benefits of one species on another that do not affect the final population density. Conditioned media experiments cannot reveal the molecular mechanisms (Dos Santos et al. 2022), but if the interactions are caused by symmetrical resource competition, we would expect an overrepresentation of reciprocal negative interactions, which we do not observe (Supplementary Figure 2CD). Although known metabolic niches would predict clear donor–acceptor patterns, our results do not fully align with these expectations, suggesting that more than simple metabolite competition and exchange underlie the observed interactions. Specifically, we expected acetate-consuming species such as *L. bacterium*, *R. intestinalis*, and *F. prausnitzii* to thrive in the spent medium of acetate producers including *A. muciniphila*, *B. adolescentis*, *B. obeum*, *B. thetaiotaomicron*, and *B. ovatus*; however, the outcomes were highly variable, indicating that additional factors, such as metabolic regulation, cross-inhibition, or signalling processes, may influence these community dynamics.

**Figure 1:**
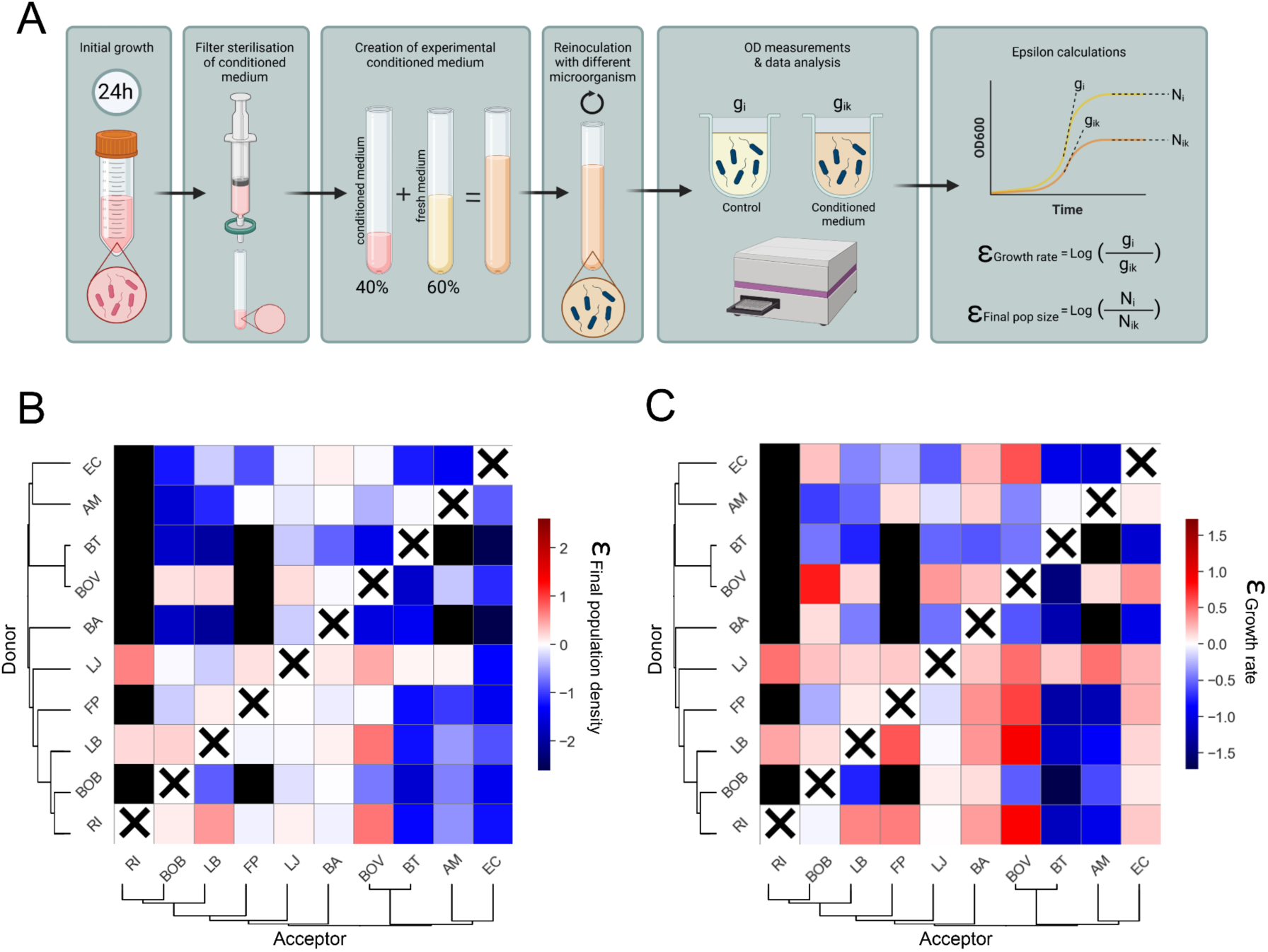
Interactions on growth rate and final population density measured by supernatant assays. **A** Experimental setup to test the invasions and calculation of the interactions. **B** Heatmap of the final population density interactions. Species are ordered by phylogenetic distance. Black boxes denote that there was no growth of the organism in the conditioned media. Species are not assessed in their own conditioned media. **C** Heatmaps of the growth rate interactions.

Grouping the organisms by their effect on other species distinguishes *L. bacterium* and *L. johnsonii* as good donor species, even excluding their effect on *R. intestinalis* (Supplementary Figure 4). We wondered if the gram type could explain species interactions. Our results show that Gram negative bacteria experience very negative interactions as acceptors and Gram positive bacteria are much less affected (Supplementary Figure 5). The phylogeny of the species also does not explain the patterns seen in the heatmaps (Figure 1), even closely related species, such as *B. ovatus and B. thetaiotaomicron*, show different effects as both donor and acceptor species. Lastly, we checked whether the monoculture growth characteristics of a species correlate with their effects as donor and acceptor species and found that species with a high monoculture final population density tend to have negative interactions, mostly as acceptor (Supplementary Figure 6). This could indicate a trade-off between monoculture growth and competitiveness with other species.

### *Roseburia intestinales* is unable to grow in most of the conditioned media

Surprisingly, three bacteria were not able to grow in conditioned media, namely *A. muciniphila* (in *B. adolescentes* and *B. thuringiensis*), *F. Praustinizii* (in the same plus *B. ovatus* and *B. obeum*) and *R. intestinalis* (in 7 out of 9 conditioned media) (Figure 1). Given that *R. intestinalis* failed to grow in many of the conditioned media samples, we considered the possibility of a batch effect. Therefore, we repeated the experiment with *R. intestinalis* as an accepted and *A. municiphila*, *B. adolescentes* and *E. coli* as donors with the same negative result. To test whether the growth inhibition was caused by toxins that kill *R. intestinalis*, we performed a spot on lawn assay. The data do not confirm killing activity of the supernatant of *A. municiphila*, *B. adolescentes* and *E. coli* (data not shown). That growth inhibition rather than lethal toxin production explains the patterns is supported by the fact that we did observe *R. intestinalis* growth in co-cultures when both species are inoculated at low densities (data not shown). Interestingly, *R. intestinalis* was positively affected by the two donors that supported growth (Figure 1). Because of the difficulty in estimating interactions when we did not observe growth, we removed *R. intestinalis* for further analyses.

### Generalised Lotka-Volterra model can predict growth in co-cultures

To test whether the growth in spent media reflects co-culture growth, we used a generalised Lotka-Volterra model to predict the equilibrium densities of co-cultures based on their interactions from the supplemented spent media (model adapted from de Vos *et al*. 2017). As the model equilibria depend solely on the final population density interactions, we refer to these as species interactions in the rest of our paper and will specify when we refer to the growth rate interactions. To validate the predictions, we grew the 36 combinations of two species and measured their final relative frequences with qPCR. Overall, the relative frequencies of the two species are well predicted by the model (Figure 2A) and clearly improve compared to predictions from monoculture growth only (Supplementary Figure 8).

**Figure 2:**
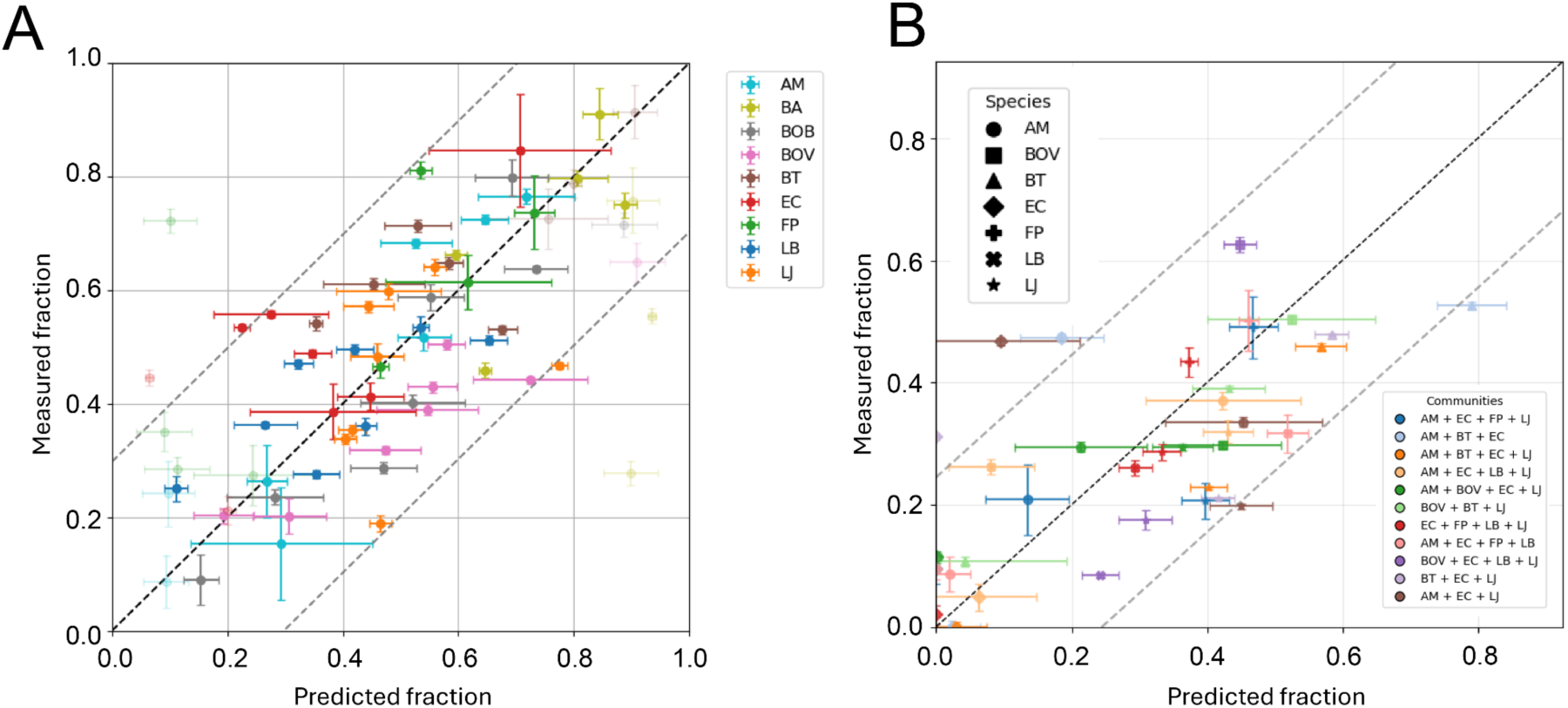
Validation of the model predictions with co-cultures of two to four bacteria. Predicted and measured species fractions are shown for co-cultures of two to four bacteria. Error bars represent standard deviations from biological replicates (measured) and 100 sensitivity analysis samples (predicted). RMSE-based dotted bands indicate the 90% confidence interval. **A:** Bi-cultures (RMSE = 0.18, Pearson r² = 0.45). **B:** Tri- and quad-cultures (RMSE = 0.14, Pearson r² = 0.50).

Exceptions to the good predictions were mainly cases where one species was not able to grow in the supplemented supernatant of another, which makes it difficult to calculate their interactions (shown opaque in Figure 2A).

We grew 11 co-cultures of 3 and 4 species for 24 hours and compared the final fraction with model predictions (Figure 2B and Supplementary Figure 7). The generalized Lotka-Volterra model with interaction terms predicts the relative abundances which cannot be predicted from monoculture growth data only (Supplementary Figure 8). Next, we estimated the possible stable communities these species could form by calculating model equilibria with all species present. We found 21.4% of the combinations to be stable with a maximum of 6 species and stable communities to have a distribution of interaction parameters that is shifted towards less negative and positive interactions (Supplementary Figure 9). Species which have more positive donor interactions on average are more likely to be found in stable communities (Supplemental Figure 10).

### Opportunistic pathogens *E. coli* and *B. ovatus* show different abilities to invade

We next wonder if these stable communities can be invaded. We first focused on the two species in our set that are reported as opportunistic pathogens, *E. coli* and *B. ovatus*. *E. coli* has a high growth rate and final population density but is negatively affected by other species, while *B. ovatus* has a lower growth rate and final population density but is positively affected by other species (Figure 3BC). When simulating invasions in stable communities, both species show different invasion patterns. *E. coli* can only invade 24.1% of the stable communities in which it is not present, mostly resulting in augmentation (Figure 3D). In contrast, *B. ovatus* can invade 97.6% of the stable communities resulting in augmentation (at low community sizes) or displacement (at high community sizes) (Figure 3D). We tested the prediction of invasion success with invasion experiments, where the stable communities are concentrated and resuspended in fresh medium plus a small amount of the invader. Relative growth of the invader was measured for 13 different communities (Figure 3E). The growth of the invader for communities with predicted invasion was significantly higher than of those where we did not predict invasion, and with a threshold of invasion as 5% of the population after 24 hours, all invasions were predicted (Figure 3F and Supplementary Figure 11).

**Figure 3:**
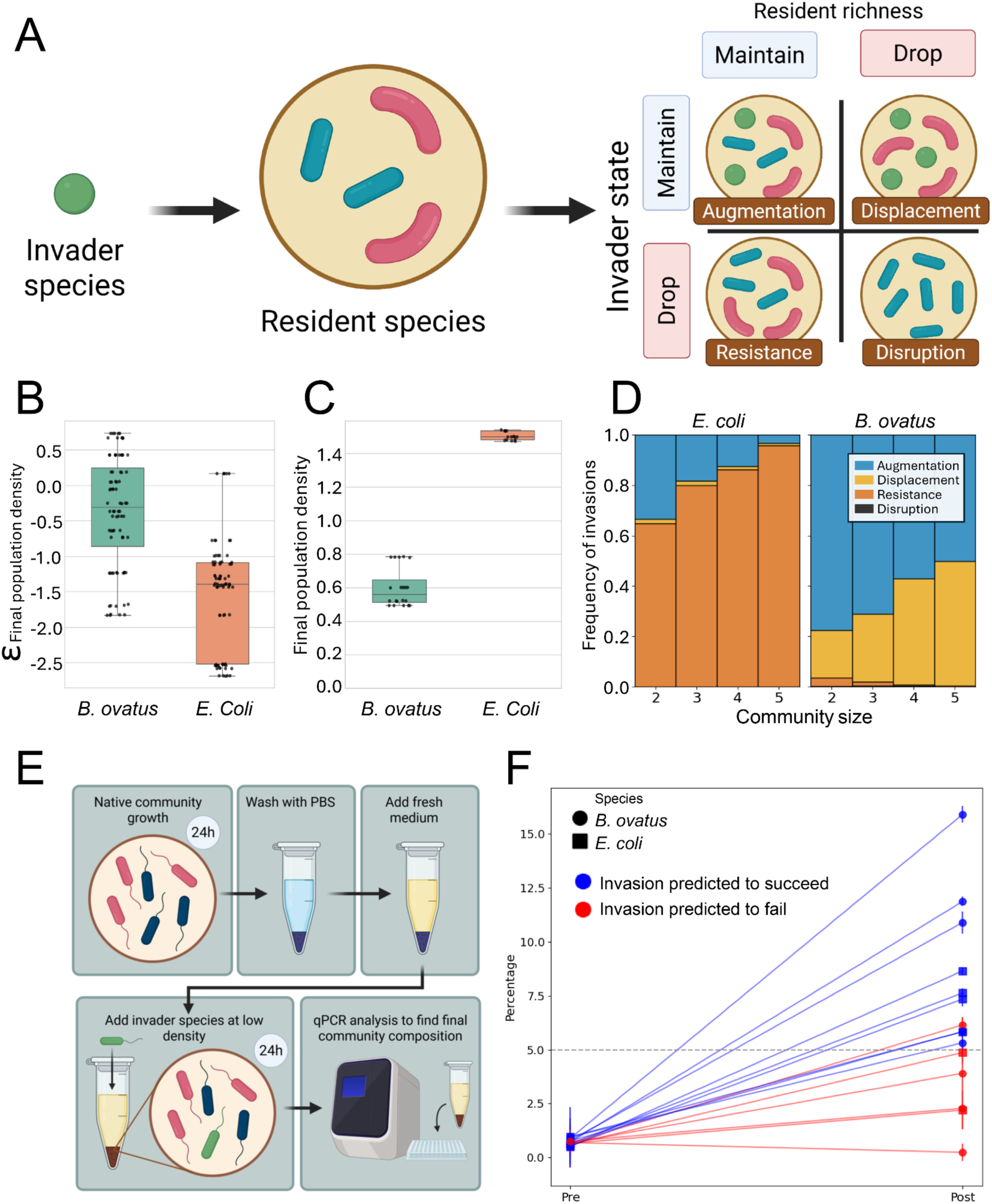
Invasion of *B. ovatus* and *E. coli* in stable communities. **A** Schematic of invasion types (adapted from (Kurkjian *et al*. 2021)). **B** Distribution of final population density interactions as acceptor with all other species (9 replicas) for *B. ovatus* and *E. coli*. **C** Final population density of *B. ovatus* and *E. coli* (9 replicas). **D** Simulation results of *E. coli* and *B. ovatus* invasions in all stable communities. **E** Schematic of the experimental procedure for testing the invasions. **F** Outcome of invasion experiments with *E. coli* (filled circles) and *B. ovatus* (filled squares) for predicted invasions (blue lines) and predicted resistant communities (red lines). The only predicted resistant community where the invader was just above the 5% threshold was repeated and then did show resistance (Supplementary Figure 11).

### Invasion can be predicted from interactions within the community and between the community and the invader

*B. ovatus* and *E. coli* exhibit distinct invasion patterns (Figure 3D), reflecting their inherent differences in growth and interaction parameters (Figures 3BC). These invasions were assessed in separate sets of stable communities, namely those that did not contain the invader. Moreover, we only assessed two species. To explore whether invader and community traits affect invasion success in an unbiased manner, we simulated species with uncorrelated characteristics (growth rates, maximum population densities, and interaction parameters) sampled from the observed distributions and tested their invasions in the full set of stable communities (see Supplementary Material S3.2). Analyzing these invasions, we observe a shift from augmentation to resistance when the community diversity increases, while displacement remains mostly constant and disruption is almost absent (Figure 4A). We scored several community and invader properties by their ability to predict invasions (Supplementary Figure 12AB). The most predictive property for the invasion is the average interaction of the community members towards the invader, where negative interactions correlate with a resistant community, and neutral and positive values result in displacement and augmentation (Figure 4B). The second most predictive property is the average interaction within the community; with an increase in the interaction coefficient displacement shifts to resistance of the community, while augmentation is hardly affected (Figure 4C). There is an interaction between the total population density and their interactions with the invader that determines invasion success (Figure 4D). At certain combinations of resident density and interaction strength, only augmentation is possible, meaning the invader can establish but cannot displace the resident community. Higher resident densities and stronger inhibitory interactions further reduce invader success, while lower densities or weaker interactions increase the likelihood of either displacement or integration. Other properties have very little impact on the invasion or are strongly correlated with the aforementioned properties (Supplementary Figure 12). In conclusion, while both negative interactions in the community and neutral to positive interactions towards the invader promote invasions, the interactions to the invader dominate, explaining the often-reported conclusion that negative interactions decrease invasion success.

**Figure 4.**
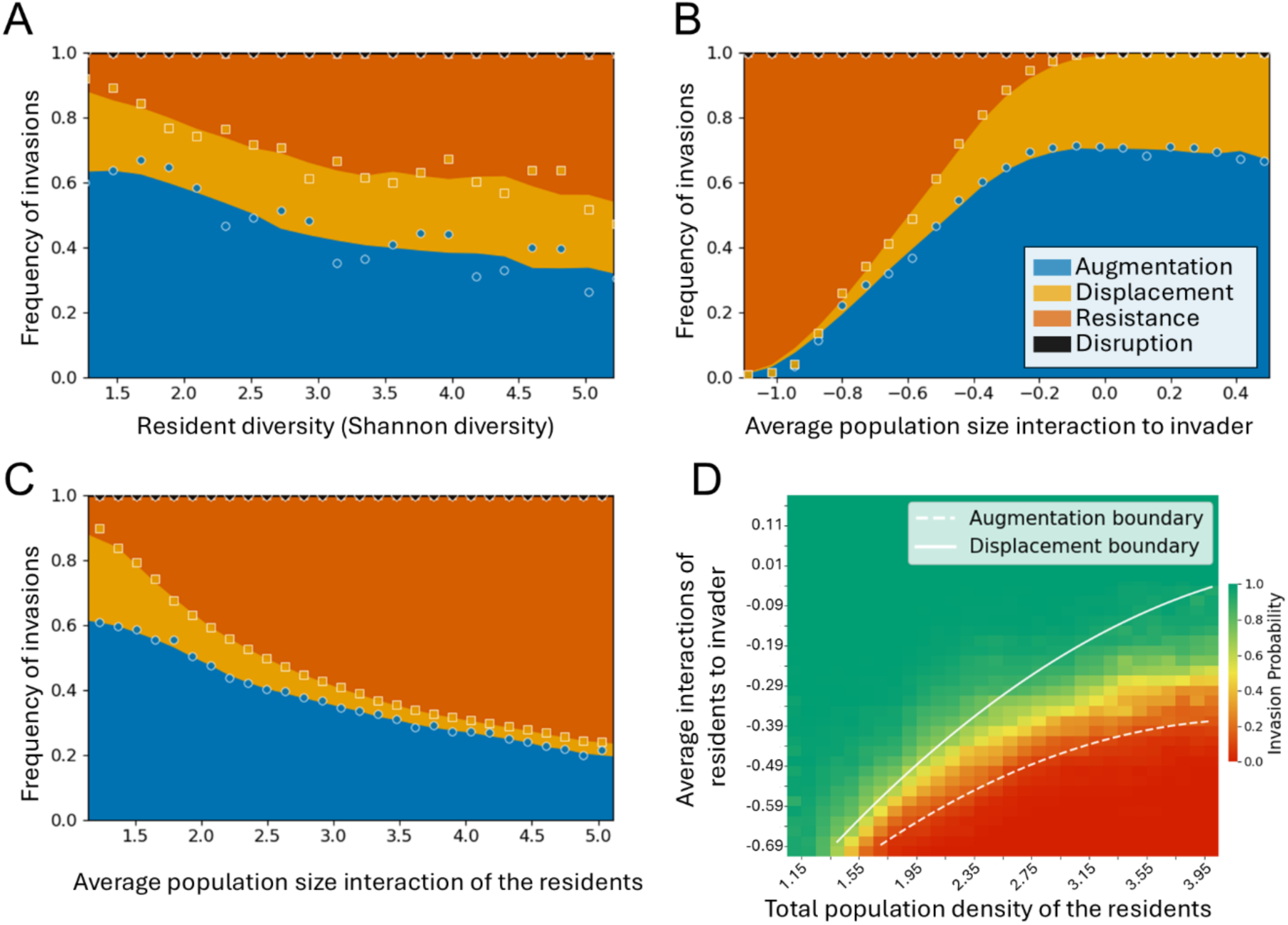
Community and invader properties affecting frequencies of different invasion types. **A** Increasing resident diversity (Shannon diversity index) decreases invasion success by decreasing the frequency of augmentation. **B** Negative interactions of the residents to the invader very strongly affect invasion success, reducing both augmentation and displacement. **C** More neutral and positive interactions between the resident species affect invasion success by almost eliminating displacement and reducing augmentation. **D** Different invasion regimes are determined by the total population dentisy of the residents and the average interactions of residents to invader. The solid and dotted lines represent the boundaries of different invasion outcomes (displacements occur only above the solid line and augmentation only above the dotted line). In the region between the two lines, any successful invader augments the resident community.

## Discussion

We have shown that environmentally mediated pairwise interactions correctly predict co-culture growth and invasion success in a SynCom of prominent gut microbiota. Commonly used species-level traits, such as Gram classification and metabolic type, failed to predict interaction outcomes, emphasizing the necessity of context-specific interaction data from microbiomes. Our results show that *E. coli* and *B. ovatus* interact differently with the other members, resulting in different invasion patterns. Community properties that predict invasions are interactions with the invader and between the community members, with diversity and total population density also exhibiting some effect, while the properties of the invaders on the resident community are of less importance. We show that the effect of the community members towards the invader dominate the invasion predictions, which is in line with previous research showing that negative interactions decrease invasion success (Hu et al. 2025; Case 1990), experiments that show that the effect of the community on the invader predict invasion outcomes (Ye et al. 2025) and that evolution of a community results in more resistance to invasion, because the member that evolves a more negative effect on the invader increases in density (Muzafar et al. 2024).

### Interactions cannot be predicted from putative metabolic interactions or Gram-type only

Ideally, interactions between species can be predicted from physiological properties such as metabolic phenotypes and Gram type. *E. coli, B. ovatus* and *B. thetaiotaomicron* are capable of assisting fermenters via cross-feeding pathways (Chia et al. 2020; Lindstad et al. 2021; Mary and Kapoor 2022), but we only observe *B. ovatus* to have positive donor interactions (Fernandez-Julia et al. 2022). We also find that *B. ovatus* has three strong positive interactions as an acceptor species, for which we do not have a metabolic hypothesis. *F. prausnitzii, L. bacterium, B. obeum* and *R. intestinalis* could be expected to be strong ‘acceptor’ species since each can consume donor-excreted metabolites present in our community (acetate for *L. bacterium* and *F. prausnitzii*; lactate for *F. prausnitzii* and *R. intestinalis*; succinate for *B. obeum*), but we we did not observe this in our conditioned media growth data (Figure 1). This indicates that the network of interactions extends beyond purely metabolic interactions: signalling via quorum sensing or other small molecules (Uhlig 2022; Markus 2023) may modulate community behaviour and obscure direct metabolite-based predictions. It is possible that some interactions are mediated by metabolic interactions that are not yet known and could play a key role in shaping the interactions we observe. Our results of the effect of Gram-type on interactions contradict earlier findings by Zandbergen *et al*. 2021, who found that Gram-positive donors tended to have stronger negative effects on other species’ growth compared to Gram-negative donors. This indicates that the effect of Gram-type is highly dependent on the specific community or environment and cannot be used as a general predictor. Additionally, while one might expect the correlation we observed between the growth rate interactions and final population density interactions (Supplementary Figure 3), this was opposite to the observations from Zandbergen *et al*. (2021). The data highlight the need for molecular physiological studies covering both primary and secondary metabolism.

### Strong response of *R. intestinalis* to other species

The inability of *R. intestinalis* to grow in conditioned medium seems to be a growth inhibitory rather than toxin mediated interaction, because we did not observe killing in a spot-on-lawn assay and did observe growth in co-cultures when both species start at low densities.

Interestingly, such a sensitive species can thrive in a diverse community as the gut microbiome. The conditioned media it did grow in, donated by *L. johnsonii* and *L. bacterium*, had a positive effect on the growth characteristics of *R. intestinalis*. We hypothesize that this beneficial relationship is mediated by lactate and acetate production of *L. jhonsonii* and *L. bacterium*, which are used as primary energy sources by *R. intestinalis*. Furthermore, both *L. jhonsonii* and *L. bacterium* can produce folate and thiamine to which *R. intestinalis* is auxotrophic, but these vitamins are present in the control medium.

### Stable community diversity and size and the effect on invasion resistance

Our model predicts co-cultures of up to four species (Figure 2). Our predictions do not seem to need higher order terms, as was the case in previous studies (D’hoe *et al*. 2018; Friedman *et al*. 2017). A diverse gut microbiome is generally seen as healthy, which can be the result of enhanced invasion resistance. We do observe this relationship (Figure 4A) and while the decrease in augmentation with community diversity agrees with the result of Zhu and Momeni 2024, we do not see the increase of displacement they report, perhaps because of the inhibitory interactions which we observe more in our community compared to their model.

### Individual versus environmental effects as predictors of invasion success

Previous studies have shown that more negative interactions lead to more invasion resistance (Case 1990; Hu et al. 2025). While we observe similar results for more negative interactions of the resident species to the invader, we find the opposite for negative interactions within the community (Fig. 4BC). These previous studies have only focussed on negative interactions and in our community we observe positive interactions, although limited. In a previous computational study with a kinetic model, the fraction of facilitative interactions (i.e. positive interactions) towards the invader have shown to increase invasion success (Kurkjian et al. 2021). Extending the analysis from Hu *et al*. (2025) to positive interactions, we observe a non-monotonic response of the invasion success to the average interactions, with maximal invasion at interactions closer to zero (Supplementary Figure 13A). Moreover, we notice that for fixed interactions with the invader, more negative interactions indeed lead to lower invasion success, similar to our findings (Supplementary Figure 13B). We can interpret our and their approach as studying idiosyncratic interactions versus environmental interactions: By decoupling the resident interactions with the interactions towards the invader we study the properties of individual differences and drawing those interactions from the same distributions (as in (Case 1990; Hu et al. 2025)), can be interpreted as the effect of the environment on interactions. While many interactions can indeed be explained by environmental effects, it remains important to separate intra resident interactions and resident-to-invader interactions, since interactions may also arise from species-specific mechanisms such as toxin production, specialized cross-feeding, or other unique physiological traits.

### Limitations of the selection of species and culture conditions

The lack of correlation between known metabolism and measured interactions could be the result of the choice of medium, which does not directly reflect the gut environment and was influenced by the fact that all species needed to be able to grow and be measured in the same medium. The lack of complex carbon sources, such as mucin, limits the possible cross-feeding interactions. We only found stable communities of up to 5 species, which is small compared to the microbial diversity observed *in vivo*. This could be because the model assumes a homogeneous environment and neglects spatial heterogeneity, host factors and differential objectives such as adhesion versus growth. Indeed, spatial heterogeneity is a driver of gut microbial assembly with distinct microbial compositions and interaction networks across intestinal regions (Mark Welch et al. 2017b; Wu et al. 2025) and host factors are known to influence colonisation and community stability (Wong et al. 2023; Paone and Cani 2020). Moreover, we only analyse 10 species, and that could limit the community size by having limited interactions profiles. Testing the interactions between a wide range of species from a specific environment and in a range of environments would paint a more complete picture of the *in vivo* situations.

In conclusion, we showed that our methods can be used to predict which species are likely to successfully engraft into existing synthetic communities. Future research can test whether these predictions can be extended to natural settings, potentially paving the way for personalized or targeted probiotics or faecal transplant donors. Enabling predictions of invasions that either enhance community resistance to pathogens or help restore disturbed microbiome states could improve the success of such treatments.

## Methods

### Bacteria and growth conditions

Bacterial strains were obtained from the DSMZ culture collection, and their identity was confirmed through 16S rRNA sequencing using universal primers, 16S forward (5’-AGAGTTTGATCCTGGCTCAG-3’); reverse (5’-ACGGCTACCTTGTTACGACTT-3’). In this study, the following bacterial strains were used: Akkermansia muciniphila (DSM 22959), Bacteroides ovatus (DSM 1896), Bacteroides thetaiotaomicron (DSM 2079), Bifidobacterium adolescentis (DSM 20083), Blautia obeum (DSM 25238), Escherichia coli (DSM 18039), Faecalibacterium prausnitzii (DSM 107838), Lachnospiraceae bacterium (DSM 24404), Lactobacillus johnsonii (DSM 10533), and Roseburia intestinalis (DSM 14610). The strains were then grown in YCFA media anaerobic conditions using an anaerobic chamber filled with anoxic gas (10% H2, 80% N2, 10% CO2) set at 37 degrees Celsius unless mentioned otherwise. The Sankey diagram for Supplementary Figure 1 was made using Sankeymatic (Bogart 2024).

### Conditioned medium preparation

We prepared conditioned medium according to De Vos *et al*. (2017). Bacteria were inoculated in YCFA medium and grown for 24 hours. After 24 hours, we measured the optical density (OD) of the culture to ensure that the bacteria had surpassed log phase growth (OD > 1). Bacteria were filtered using a 0.2-micron filter to obtain the spent medium. We then prepared conditioned medium by mixing 60% fresh medium with 40% of spent medium.

### Growth of bacteria in conditioned medium

For the growth experiments, bacteria were pre-cultured for 16 hours or until they reached logarithmic phase. The bacteria were then diluted to an OD of 0.05 in conditioned medium. All 10 bacteria were cultured in all other 9 conditioned media, resulting in a total of 90 combinations. The growth of the bacteria was monitored using a Byonoy plate reader at a wavelength of 595 nm for 24 hours in an anaerobic workstation (Whitley A35 Workstation, Don Whitley Scientific) under anaerobic conditions (10% H2, 10% CO2, 80% N2) at 37 degrees Celsius.

### Co-culture growth experiments

To grow the co-cultures, the bacteria were first individually pre-cultured for 16 hours or until they reached logarithmic phase. Using OD-CFU correlations, the bacteria were combined using an equal number of CFU’s and subsequently diluted to a final combined OD of 0.05. All co-cultures were grown for 24 hours in 8mL flasks. After 24 hours, the OD was measured and the samples were prepared for downstream applications.

### Invasion assay

Initial consortia were cultivated for 24 hours starting at an OD600 of 0.05. Simultaneously, a preculture of the invading species was grown under identical conditions. After 24 hours, the OD600 of the initial consortia cultures was measured. The bacterial cultures were centrifuged at 13000 x g for 5 minutes, and the supernatant was discarded. The cell pellets were resuspended in fresh growth medium. The invading species was washed twice with PBS, and its OD600 was measured. The washed invading species was then introduced into the resuspended initial consortia at a final OD600 of 0.05. Followed by an additional 24-hour incubation period. Total DNA was isolated from the cultures and then used for the qPCR assay.

### DNA isolation

DNA isolation was performed using a modified Phenol-Chloroform method. Co-cultures were grown as described above and centrifuged at max speed to pellet the cells. The pellet was resuspended in elusion buffer (10 mM Tris-HCl, pH 8.5). The suspension was then mixed with an equal volume of phenol:chloroform:isoamyl alcohol (25:24:1) to separate the DNA from other cell components. The mixture was centrifuged, and the DNA-containing aqueous layer was transferred to a new container. The DNA was subsequently precipitated by adding ethanol and ammonium acetate to a final concentration of 0.75M. The DNA was collected by centrifugation, washed with 70% ethanol, and dissolved in elution buffer. The purified DNA was then quantified using spectrophotometry and stored at −20 degrees Celsius for downstream applications.

### qPCR analysis of community composition

Primers targeting the bacterial species of interest were designed using primer3plus (https://www.bioinformatics.nl/cgi-bin/primer3plus/primer3plus.cgi) to amplify either a specific region of the bacterial 16S rRNA gene or a specific gene for the focal species and yield a PCR product of approximately 200 base pairs with a delta of 50 base pairs (Supplementary Table S1). The designed primers were tested against all the other bacterial species used in this study to verify their specificity and exclude any potential false positives. PCR amplification was performed and followed by gel electrophoresis to confirm the expected product size and absence of nonspecific amplification. The SYBR green method was employed for quantitative PCR (qPCR) analysis. The qPCR reactions were performed in a total volume of 25 μL, containing 12.5 μL of SYBR green master mix, 1 μL of each forward and reverse primer (10 μM), 2 μL of template DNA, and 8.5 μL of nuclease-free water.

Negative controls (no template DNA) were included in each run to monitor potential contamination. qPCR amplification was carried out using the Applied Biosystems 7300 Real-Time PCR with the following thermal cycling conditions: an initial denaturation step at 95°C for 5 minutes, followed by 40 cycles of denaturation at 95°C for 30 seconds, annealing at 60°C for 30 seconds, and extension at 72°C for 30 seconds. The fluorescence signal was measured at the end of each extension step. The quantification cycle (Cq) values were recorded in triplicate for each sample, and averages were used. Standard curves were generated using known concentrations of bacterial DNA containing the target sequence. The Cq values of the samples were compared to the standard curve to determine the initial bacterial load in each sample. Positive controls with known concentrations of the bacterial species were included to assess the efficiency and sensitivity of the qPCR assay. See Supplementary Material S1.3 for the calculation of the errors.

### Modelling and data analysis

All data analysis was done in rStudio using R version 4.3.1 and Python version 3.11.4. Growth parameters were extracted through the growth rates package (Petzoldt 2022). Co-culture predictions were made using the generalized Lotka–Volterra equations incorporating a logistic growth formulation (de Vos et al. 2017), with minor modifications (see Supplementary Material S2 for modifications and motivation). Co-culture predictions were calculated from the equilibrium points of the sets of differential equations (Supplementary Material S2.2). Error bars on the predictions were calculated with simulations with parameters drawn from a normal distribution using the standard deviations calculated from the measurement error (Supplementary Material S1.2). To evaluate the quality of the fit between the predicted and measured data, the root-mean-square error (RMSE) was computed as a measure of the average deviation between predicted and measured values. In addition, the strength of the linear relationship between measured and predicted values was quantified using the Pearson squared correlation coefficient (r²). The Pearson correlation coefficient was obtained using the linregress function from the scipy.stats Python library. For the invasion assays, the model was simulated for 48 hours and a threshold equal to the inoculum value was used to determine species persistence (see Supplementary Material S3.1).

## Data availability

All data and code are available in the GitHub repository: https://github.com/WortelLab/SynCom

## Acknowledgements

We acknowledge Gertien Smits for discussions at the initialisation of the project and Tobias Köhler and Richard de Boer for help with the qPCR experiments. Supported by University funds dedicated to the Research Priority Area Systems Biology of Host Microbiome Interactions.

## Notes

https://github.com/WortelLab/SynCom

### Competing Interest Statement

The authors have declared no competing interest.

### Summary of Updates

Data is reanalysed, figures are updates, supplementary figures and material are added. Some experiments are added. Sampling of invaders and figure 4 are new. Extra author is added. Paper is rewritten.

