## Supplemental Material for "Environmentally mediated interactions predict community assembly and invasion success in a gut microbiota SynCom"

#### Supplementary Figures

**Supplementary Figure 1: The inclusion of bacterial strains.** **3**
**Supplementary Figure 2: Classification and prediction of pairwise interactions.** **4**
**Supplementary Figure 3: Correlation between bacterial growth and yield interaction effects.** **5**
**Supplementary Figure 4: donor and acceptor interaction first principal component.** **6**
**Supplementary Figure 5: Mirror distributions of interaction strengths between Gram-positive and Gram-negative bacteria.** **7**
**Supplementary Figure 6: Relationship between mono-culture data and interaction effects.** **8**
**Supplementary Figure 7: Comparison of in vitro and in silico community compositions for tri- and quad-species communities.** **9**
**Supplementary Figure 8. Model predictions without interaction parameters deviate substantially from experimental observations in co-cultures of two to four species.** **10**
**Supplementary Figure 9: Properties of stable communities.** **12**
**Supplementary Figure 10: Effect of average interactions on the persistence of bacterial species across community sizes.** **13**
**Supplementary Figure 11. Properties of the communities used in the invasion experiments.** **14**
**Supplementary Figure 12. Properties of communities and invaders most predictive of invasion outcomes.** **15**
**Supplementary Figure 13: Comparison with the result from Hu et al (2025).** **16**
**Supplementary Table 1: Strains used in the study with qPCR primers** **17**

### Supplementary Material

**S1 Parameters and Sensitivity Analysis** 17
- **S1.1 Model Parameters** 17
- **S1.2 Parameter Sampling via Sensitivity Analysis** 17
- **S1.3 Error Propagation for qPCR-Derived Concentrations** 17
- **S1.4 Uncertainty of Interaction Parameters** 18

**S2 Theoretical Model of Microbiota Interactions** 19
- **S2.1 Comparison with an ecological variant** 20
- **S2.2 Computation of community endpoints** 20

**S3 Invasion Assay Using the Theoretical Model** 20
- **S3.1 Simulating invasion into previously stabilized communities** 20
- **S3.2 Generation of random invader species** 21

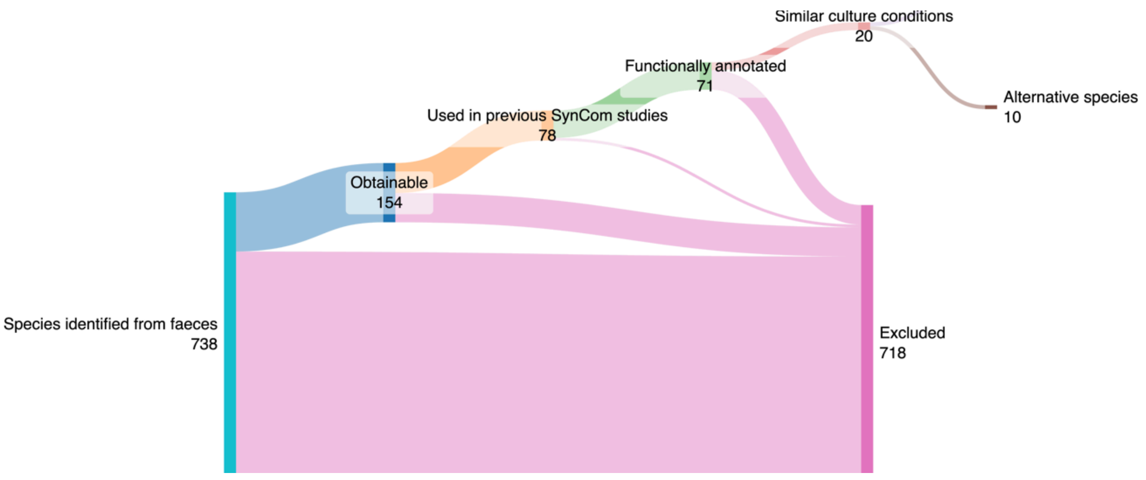


**Supplementary Figure 1: The inclusion of bacterial strains.**

Starting with 738 bacteria, we used different filtering steps to reduce the number of strains to be used in our experimental setup down to 10. We started with bacterial strains that are known to be present in the western human gut. From that, we selected strains that have been previously identified as being associated with beneficial effects on human health and strains that have been identified as being associated with disease. Furthermore, we made sure all strains we wanted to use were obtainable. Additionally, to increase the likelihood of co-growth in a laboratory setting we selected species that were previously used in SynComs. In summary, the species selected all have clearly defined functions within the microbiome, have known sequences of at least their 16S region, and are all culturable under similar anaerobic conditions in a lab environment. Then we selected a set of ten species that were representative of gut composition on a phylum level, making sure all large categories of functions were covered. See Table S1 for the list of species. Ten bacteria were left that can be used as an incidental backup, but were not used in this study.

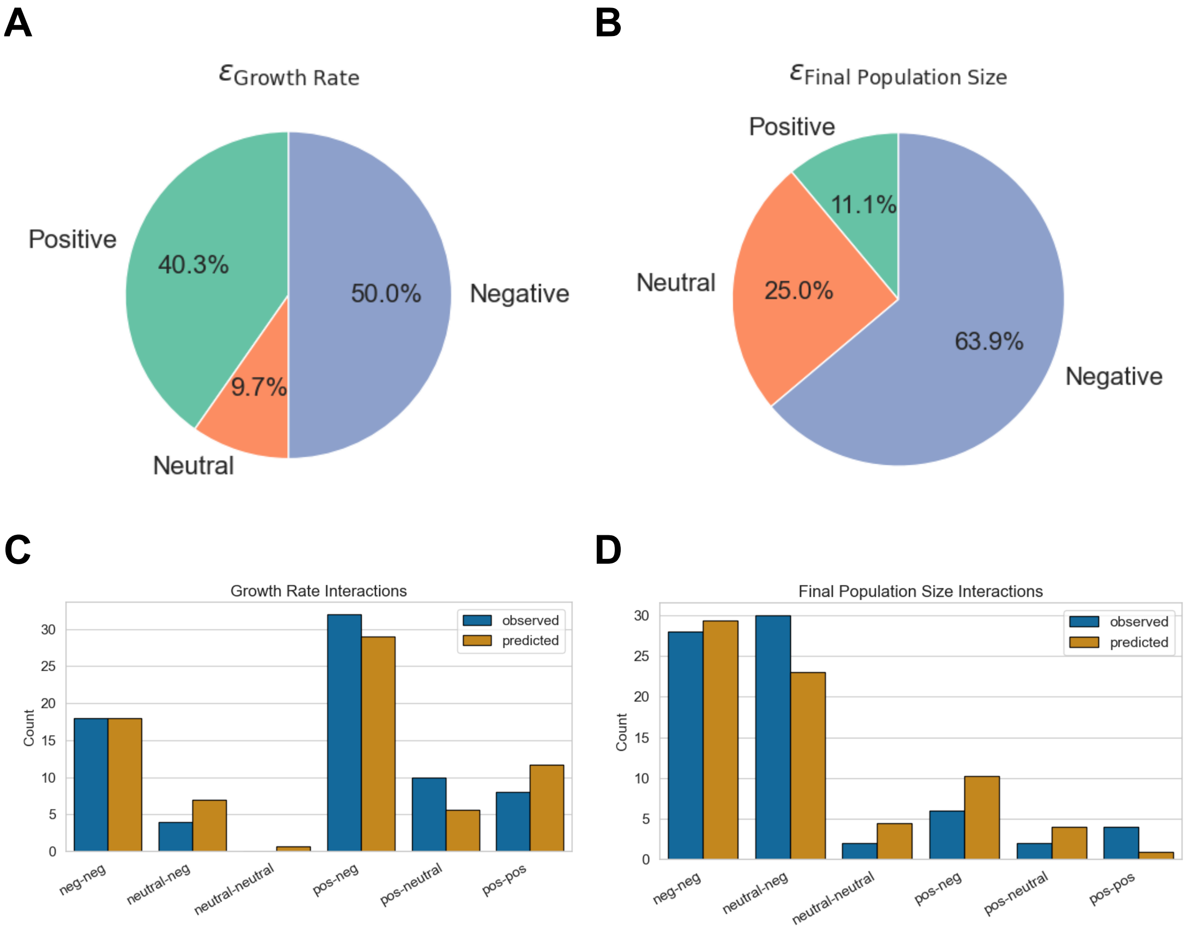


**Supplementary Figure 2: Classification and prediction of pairwise interactions.**

The cutoff for neutral interactions was set between -0.1 and +0.1. RI is excluded from this analysis. **A:** Percentage of different types of growth rate interactions. **B:** Percentage of different types of final population size interactions. **C:** Observed vs predicted pairwise growth rate interactions. **D:** Observed vs predicted pairwise final population size interactions. None of the observed versus predicted distributions were significantly different according to a binomial test (minimum total count 5).

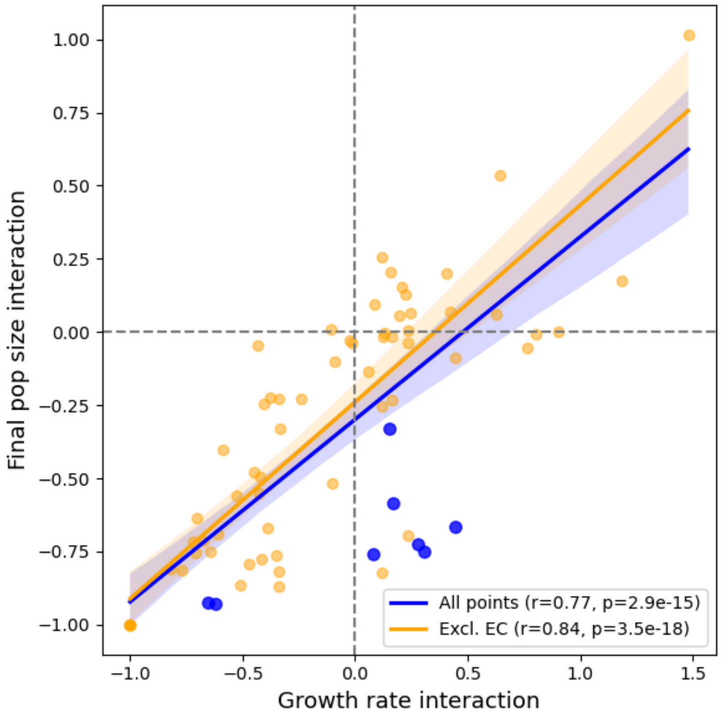


**Supplementary Figure 3: Correlation between bacterial growth and yield interaction effects.**

Each point represents a pairwise interaction between bacterial species, showing the relationship between mean growth effects and mean final population size effects. The blue points correspond to *E. coli* (EC) interactions, while orange points represent all other species. Regression lines depict correlations calculated across all data (blue) and excluding *E. coli* (orange). A positive association indicates that species promoting higher growth also tend to enhance final population size.

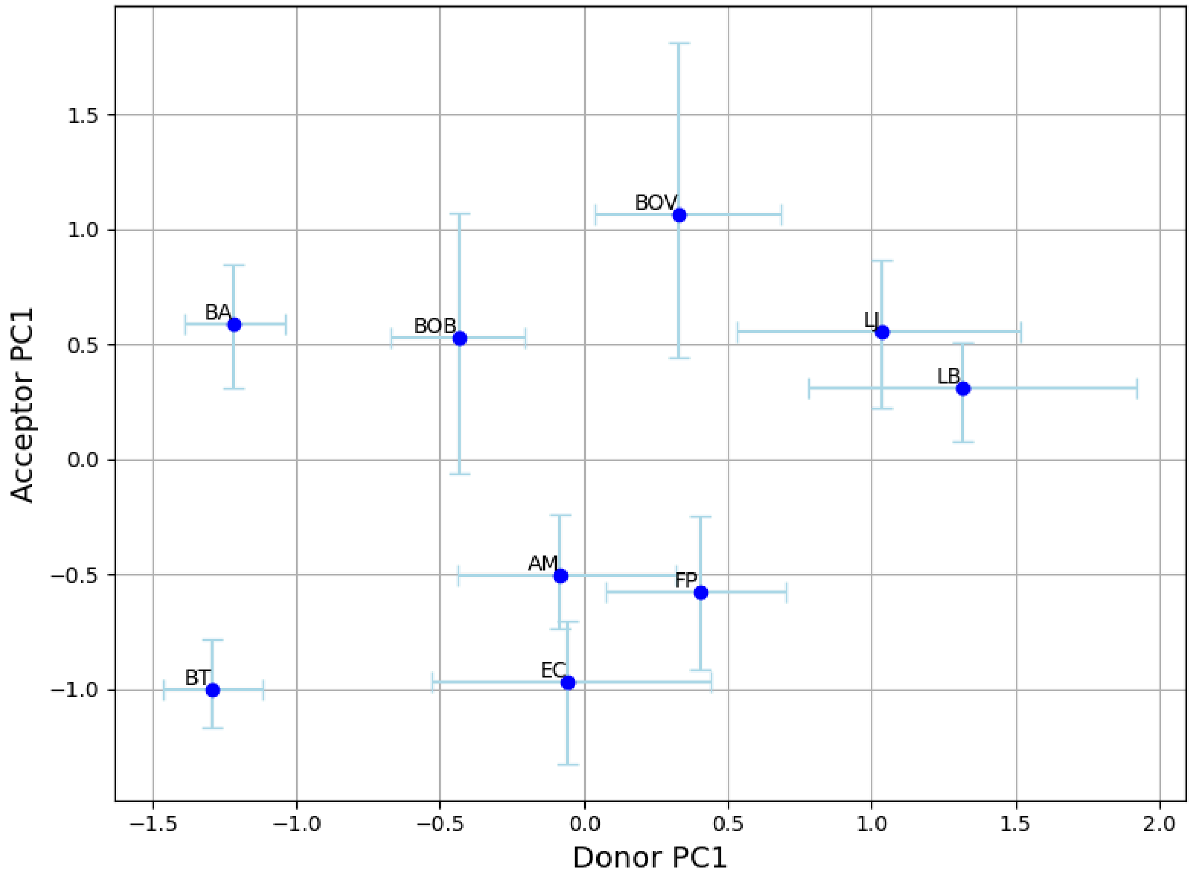


**Supplementary Figure 4: donor and acceptor interaction first principal component.**

Each point represents a bacterial species, plotted according to its mean PC1 score from the donor (x-axis) and acceptor(y-axis) perspectives. Error bars denote 95% confidence intervals obtained from 10,000 bootstrap samples. The positive association between donor and acceptor PC1 scores indicates that species contributing strongly to others' final population size (donors) also tend to benefit more from community interactions (acceptors). Benefit PC1 explains on average 45% (95% CI: 39%–51%) of the variance, while Acceptor PC1 explains 54% (95% CI: 46%–61%) of the variance. See table S1 for species abbreviations.

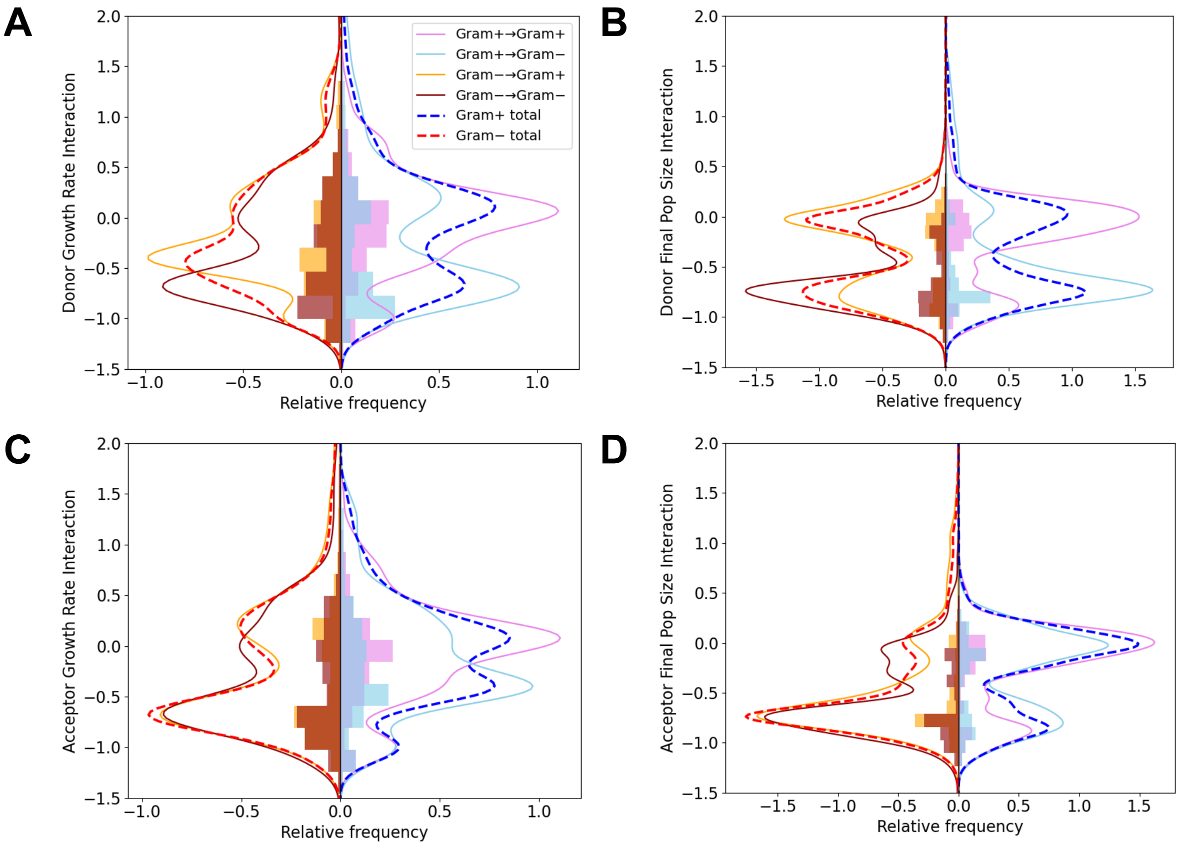


**Supplementary Figure 5: Mirror distributions of interaction strengths between Gram-positive and Gram-negative bacteria.**

For each interaction category (GramPositive→GramPositive, GramPositive→GramNegative, GramNegative→GramPositive, GramNegative→GramNegative), we bootstrap 100 values per observation from a Normal distribution parameterized by the experimental mean and standard deviation, then plot relative-frequency histograms and kernel density estimates (KDE). The horizontal axis shows relative frequency/KDE (positive for Gram-positive focal, negative for Gram-negative focal); the vertical axis is the interaction value. Dashed curves summarize the total KDE for Gram-positive focal (blue) and Gram-negative focal (red). **A:** donor growth rate interactions, **B:** donor final population size interactions, **C:** acceptor growth rate interactions, **D:** acceptor final population size interactions.

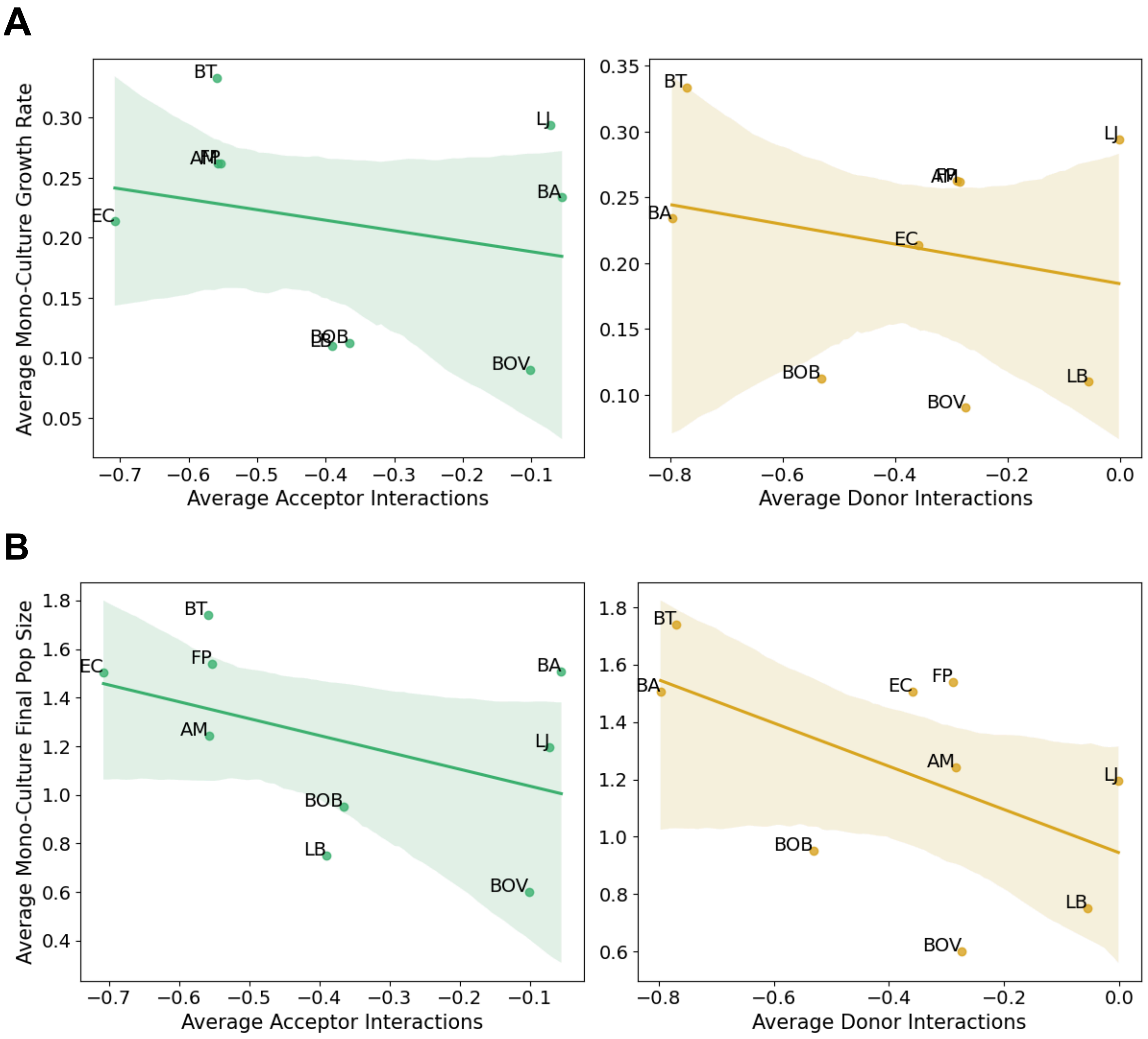


**Supplementary Figure 6: Relationship between mono-culture data and interaction effects.**

Each point represents a bacterial species, showing the association between its mean monoculture data and the average magnitude of its interactions as a community acceptor (left) or donor (right). Regression lines indicate overall trends, suggesting that higher final population size and faster-growing species in mono-culture tend to interact less with others. **A** Relationship between monoculture growth rate and average final population size interaction effects. **B** Relationship between monoculture final population size and average final population size interaction effects. See table S1 for species abbreviations.

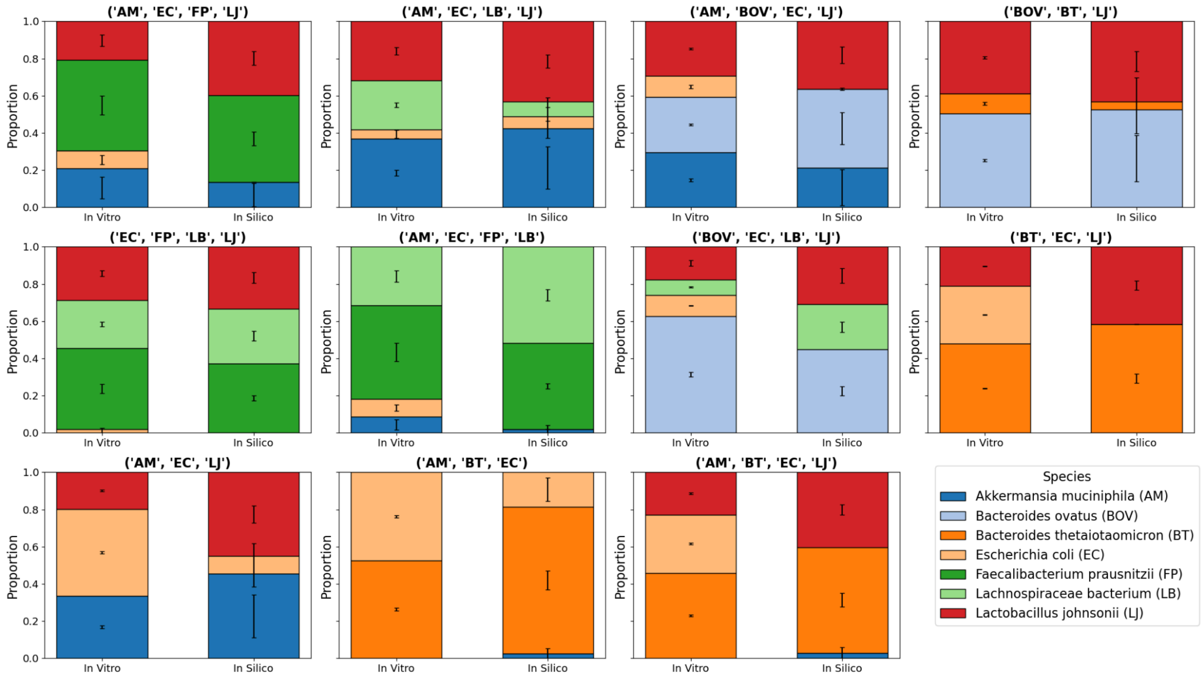


**Supplementary Figure 7: Comparison of in vitro and in silico community compositions for tri- and quad-species communities.**

Stacked bar plots show the relative abundance of each species within experimentally observed (*in vitro*, left) and model-predicted (*in silico*, right) communities. Each subplot represents a unique tri- or quad-species culture, with colors denoting individual species. Error bars indicate the standard deviation across the experimental replicates (left) or across the model result from the sensitivity analysis (right).

**A**

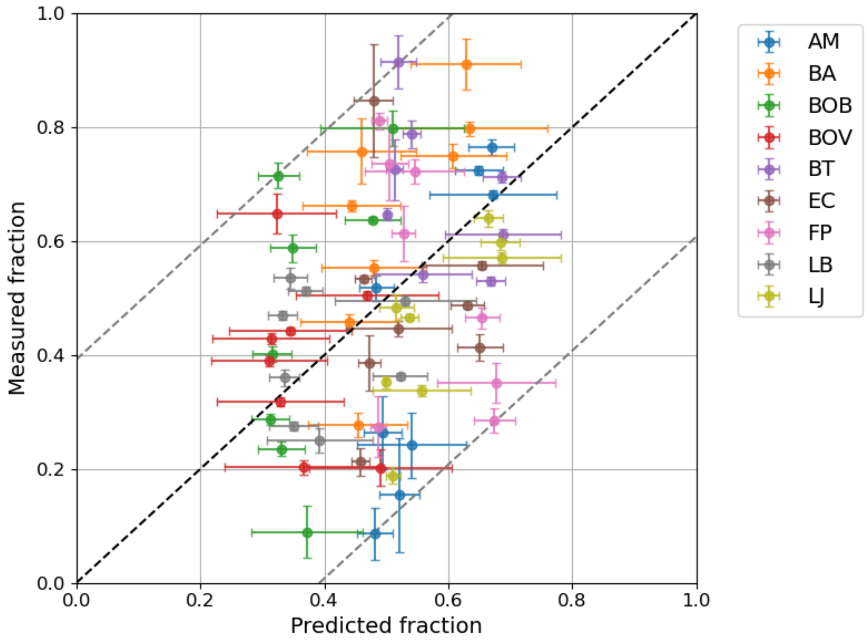


B

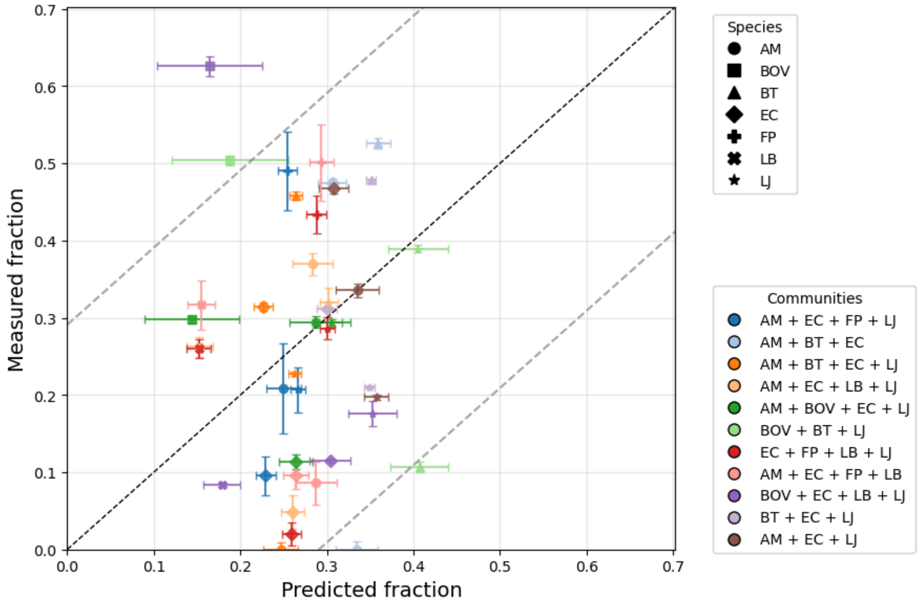


**Supplementary Figure 8. Model predictions without interaction parameters deviate substantially from experimental observations in co-cultures of two to four species.**

Predicted and measured species fractions are shown for co-cultures containing two to four bacterial species using a model in which all interaction terms are set to zero with a time simulation of 24 hours. Error bars indicate standard deviations across biological replicates (measured) and across 100 sensitivity-analysis samples (predicted). Dotted bands represent 90% confidence intervals based on RMSE. A: Two-species co-cultures (RMSE = 0.2, Pearson $r^2 = 0.10$). B: Three- and four-species co-cultures (RMSE = 0.18, Pearson $r^2 = 0.00$). See table S1 for species abbreviations.

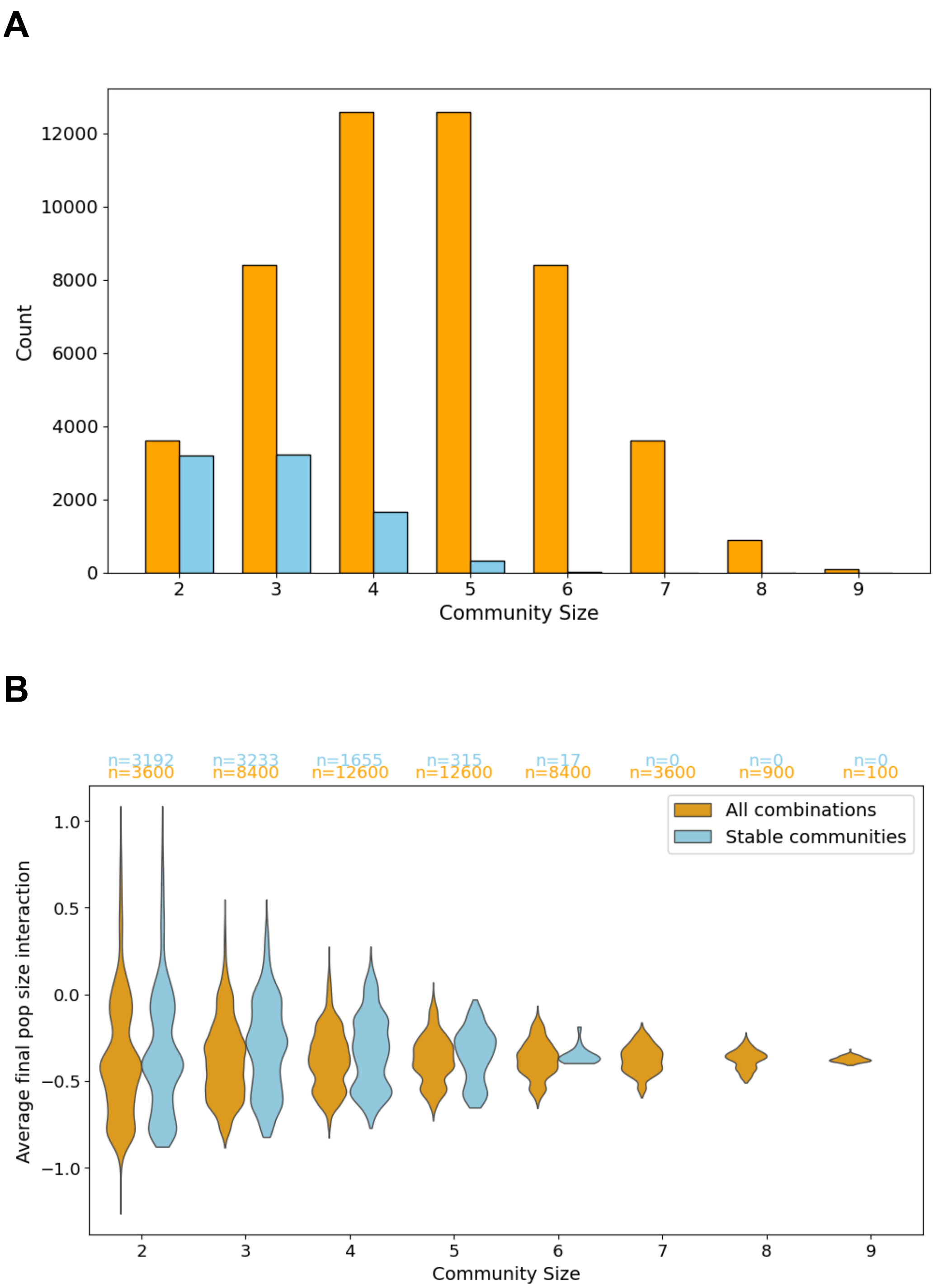


**Supplementary Figure 9: Properties of stable communities.**

**A** Total combinations and stable communities of different community sizes. **B** Comparison of interaction strengths between all community combinations and stable communities across community sizes.Violin plots illustrate the distribution of the average final population size resident interaction values for all simulated combinations (orange) and the stable communities (blue). Each pair of violins corresponds to a specific community size, highlighting how stable communities tend to exhibit slightly higher positive interactions for larger community size. Sample sizes for each group are indicated above the violins.

**A**

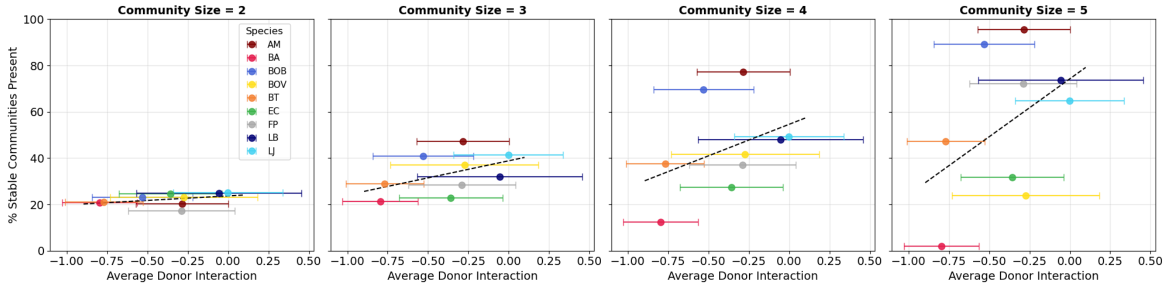


**B**

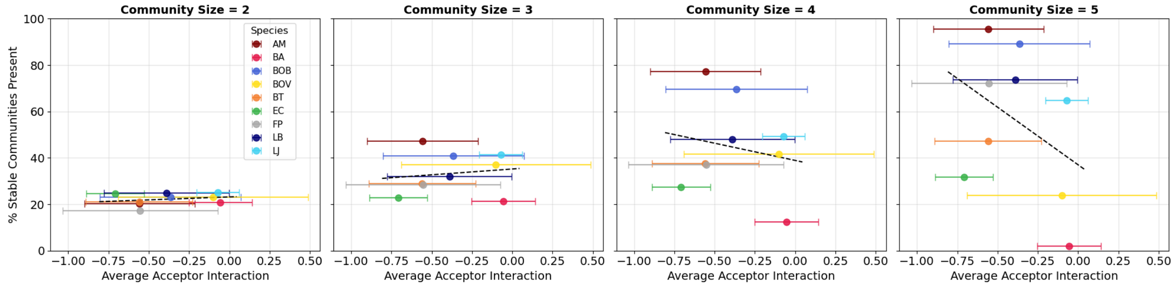


**Supplementary Figure 10: Effect of average interactions on the persistence of bacterial species across community sizes.**

Each panel represents a different community size, showing the relationship between a species' average donor or acceptor final population size interaction strength (x-axis) and its frequency of presence in stable communities (y-axis). Points denote species means ± standard deviation, and dashed lines indicate linear regressions. A positive correlation suggests that species exhibiting stronger positive effects on others (right) or benefit more from others (right) are more likely to persist across increasingly complex community assemblies. **A:** Effect of average donor interactions on the persistence of bacterial species across community sizes. **B:** Effect of average acceptor interactions on the persistence of bacterial species across community sizes.

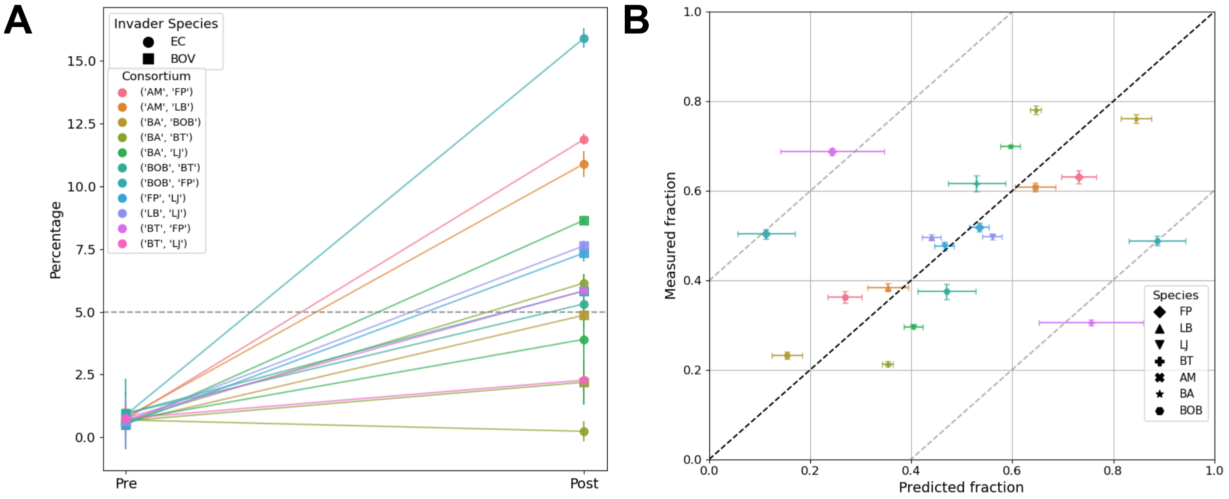


**Supplementary Figure 11. Properties of the communities used in the invasion experiments.**

**A:** Outcomes of invasion experiments with *E. coli* (filled circles) and *B. ovatus* (filled squares) across bi-culture stable communities. Colors indicate different communities. **B:** Predicted and measured species fractions before invasion. Community colors match those in panel A. See table S1 for species abbreviations.

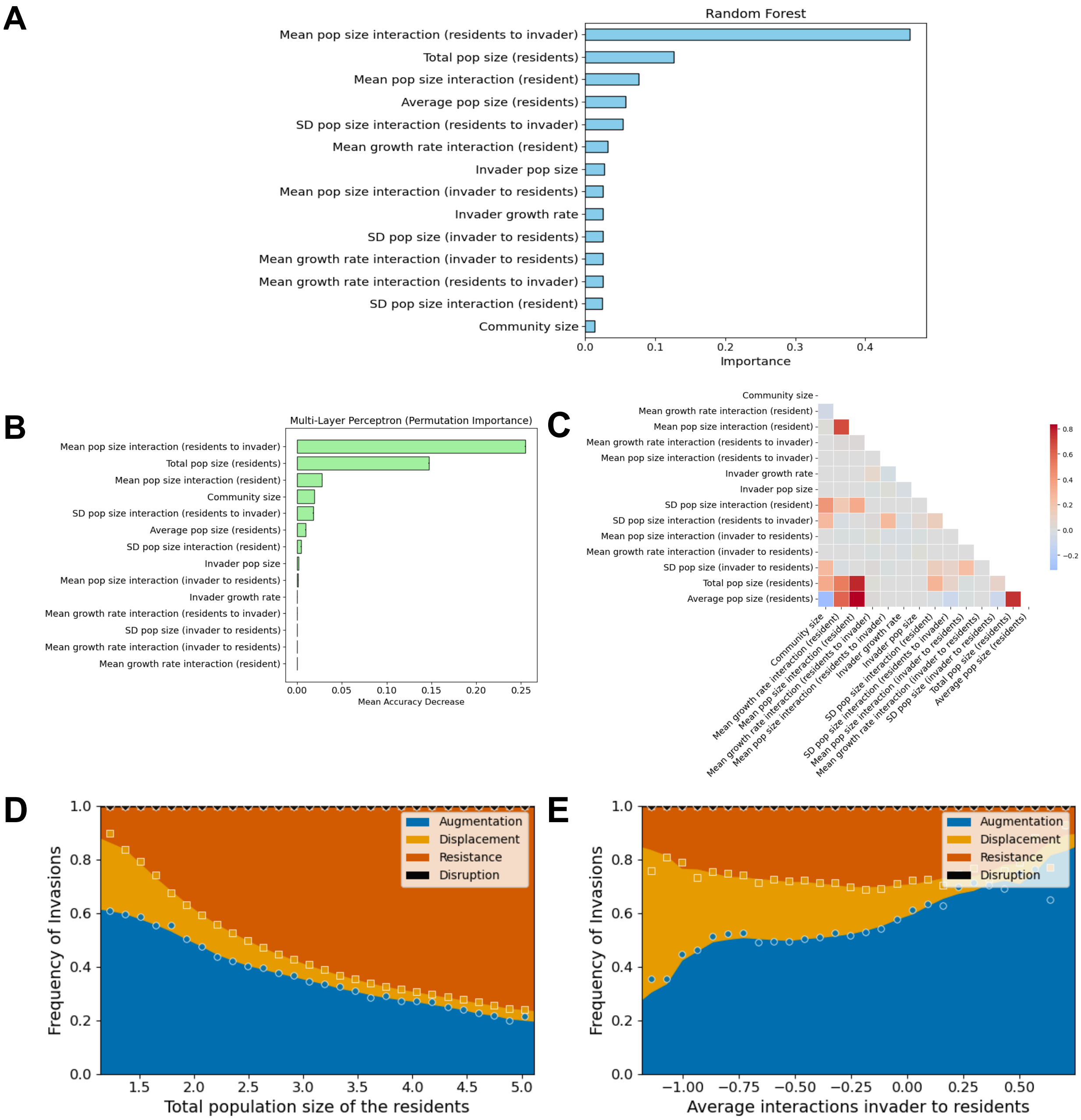


**Supplementary Figure 12. Properties of communities and invaders most predictive of invasion outcomes.**

**A:** Feature importance from the Random Forest model (accuracy = 0.97). Features are ordered by importance; higher values indicate stronger influence on predicted invasion success. **B:** Feature sensitivity for the MLP model (accuracy = 0.92) using permutation importance. Mean decrease in model accuracy is shown for each feature when shuffled, with error bars representing standard deviation across permutations.**C:** Pairwise correlations among key features describing community composition, resident and invader properties, and interaction effects. Colors indicate correlation strength and direction (red = positive, blue = negative); only the lower triangle is displayed for clarity. **D:** The effect of average total population size of residents on invasion frequency is a strong predictor but highly correlated with the effect of resident-resident population interactions. **E:** The effect of average population interactions from invader to residents is a poor predictor of invasion frequency.

**A**

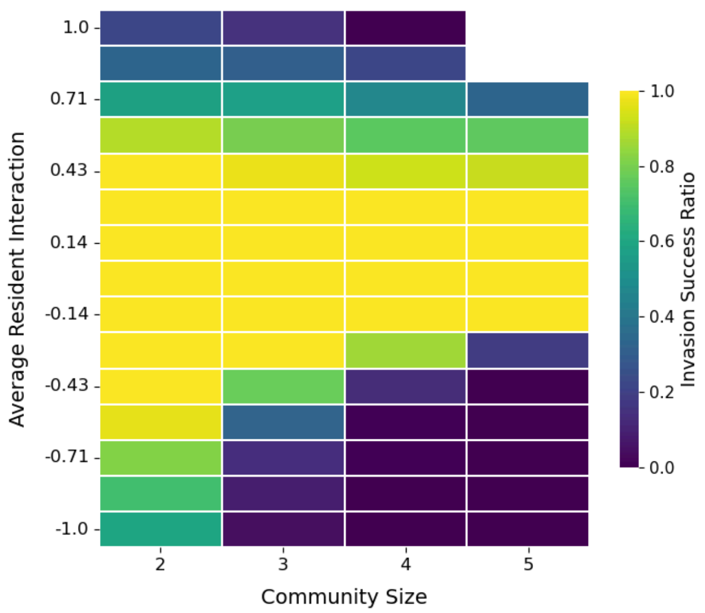


**B**

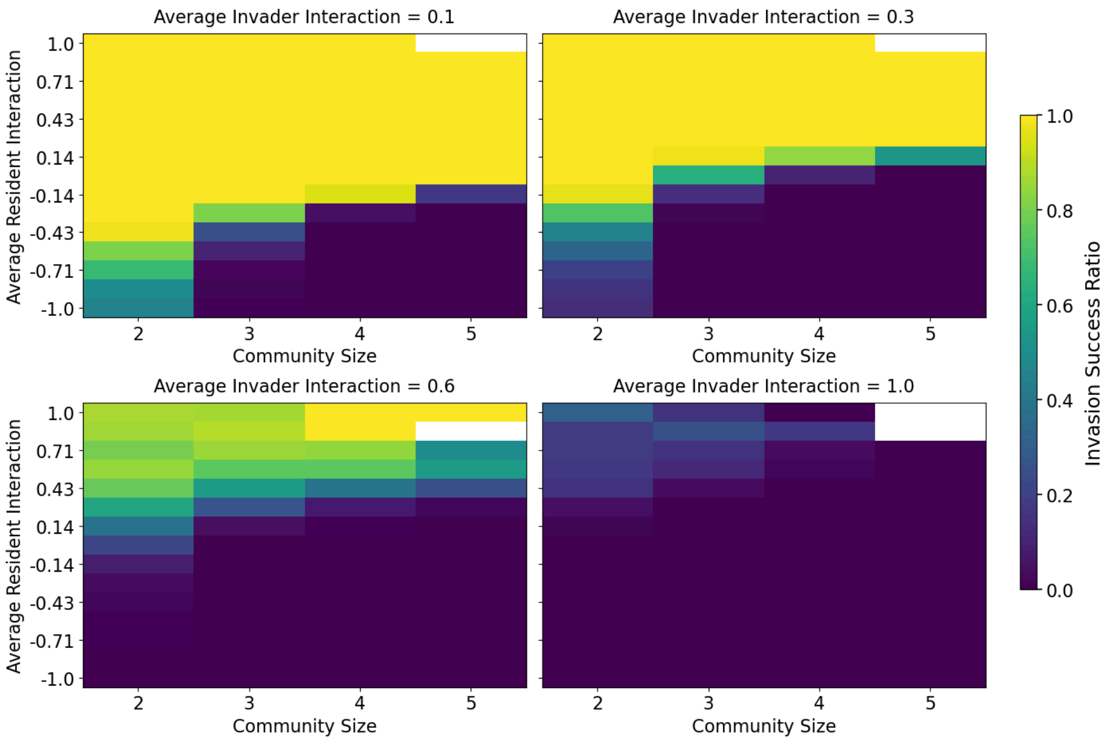


**Supplementary Figure 13: Comparison with the result from Hu et al (2025).**

Invasion success across stable communities of varying size and resident interaction strength using a Generalized Lotka–Volterra model. Interactions are drawn following the approach of Hu *et al.* (2025), and no dispersal is considered. **A:** Invader interactions are drawn as in Hu *et al.* (2025). **B:** Invader interactions are independent of the resident community.

**Supplementary Table 1: Strains used in the study with qPCR primers**

| Strain | Primer | Sequence (5′-3′) | Product size (bp) |
|---|---|---|---|
| *Akkermansia muciniphila* (AM) DSM 22959 | Fw | GACATGCAAGTCGAACGAGA | 241 |
| | Rev | CATCCTCTCAGACCGGCTAC | |
| *Bacteroides ovatus* (BOV) DSM 1896 | Fw | AACTCCGGAATAGCCTTTCG | 208 |
| | Rev | CGTAGGAGTTTGGACCGTGT | |
| *Bacteroides thetaiotaomicron* (BT) DSM 2079 | Fw | CGTTCCATTAGGCAGTTGGT | 193 |
| | Rev | CAACCCATAGGGCAGTCATC | |
| *Bifidobacterium adolescentis* (BA) DSM 20083 | Fw | AACCTTACCTGGGCTTGACA | 238 |
| | Rev | CGTAAGGGGCATGATGATCT | |
| *Blautia obeum* (BOB) DSM 25238 | Fw | CCGCGTGAAGGAAGAAGTAT | 247 |
| | Rev | CTTACCTCTCCGGCACTCAA | |
| *Escherichia coli* (EC) DSM 18039 | Fw | CAACTGAAGCGTCAGCAAAG | 225 |
| | Rev | GCTGGAAGTGCAGTCAACAA | |
| *Faecalibacterium prausnitzii* (FP) DSM 107838 | Fw | GTGCTGGAAGCCGACACC | 246 |
| | Rev | CGCAGCGTCCAGAAACTT | |
| *Lachnospiraceae bacterium* (LB) DSM 24404 | Fw | CATCCAGAGTCCCCTTCAAA | 214 |
| | Rev | CTGGGGCTTATGAACGTTGT | |
| *Lactobacillus johnsonii* (LJ) DSM 10533 | Fw | GCGAGCTTGCCTAGATGATT | 244 |
| | Rev | ATCGCCTTGGTAAGCCATTA | |
| *Roseburia intestinalis* (RI) DSM 14610 | Fw | TGGACGATTACTGACGCTGA | 244 |
| | Rev | TAAGGTTCTTCGCGTTGCTT | |

### Supplementary Material

#### S1 Parameters and Sensitivity Analysis

##### S1.1 Model Parameters

The parameters used to parameterize the theoretical model are defined as follows:

- Intrinsic growth rate of species $i$, $g_i$: estimated from monoculture assays in fresh medium.
- Final population density of species $i$, $\hat{X}_i$: estimated from monoculture assays in fresh medium.
- Growth-rate interaction matrix $A = (a_{ij})$: estimated from monocultures grown in fresh and in conditioned media. Each entry $a_{ij}$ quantifies the multiplicative change in the growth rate of species $i$ when cultured in conditioned medium produced by species $j$, relative to its growth rate in fresh medium.
- Final-density interaction matrix $B = (b_{ij})$: estimated from monocultures grown in fresh and in conditioned media. Each entry $b_{ij}$ quantifies the multiplicative change in the final density of species $i$ when cultured in conditioned medium produced by species $j$, relative to its final density in fresh medium.

##### S1.2 Parameter Sampling via Sensitivity Analysis

All simulations—both for assessing stable community states and for testing invasions—were performed using parameters sampled from distributions that reflect experimental uncertainty. Specifically, for each model run we draw

$$g_i \sim \mathcal{N}(\bar{g}_i,\, \sigma^2_{g,i}), \qquad \hat{X}_i \sim \mathcal{N}(\bar{\hat{X}}_i,\, \sigma^2_{\hat{X}_i}), \qquad a_{ij} \sim \mathcal{N}(\bar{a}_{ij},\, \sigma^2_{a,ij}), \qquad b_{ij} \sim \mathcal{N}(\bar{b}_{ij},\, \sigma^2_{b,ij}),$$

where $\bar{\cdot}$ and $\sigma_\cdot$ denote the empirical mean and standard deviation obtained from experimental measurements. This sampling scheme captures natural variability and measurement noise, ensuring that community assembly and invasion dynamics are evaluated across a broad set of plausible parameter combinations.

##### S1.3 Error Propagation for qPCR-Derived Concentrations

Because the experimental error dominates technical variation, only biological replicates at the cycle threshold (Ct) cutoff are considered; technical replicates are not included.

Let the linear calibration from Ct to concentration be

$$Y = aX + b, \tag{1}$$

where $X$ is the Ct value and $Y$ the estimated bacterial concentration. If the uncertainty is reported on Ct (standard deviation $\sigma_X$), the concentration uncertainty is obtained by linear propagation:

$$\sigma_Y = |a|\,\sigma_X.$$

Conversely, if the uncertainty is reported on concentration, the corresponding Ct uncertainty is

$$\sigma_X = \frac{\sigma_Y}{|a|}.$$

For two-species mixtures, the fraction of species $Y$ is

$$p_Y = \frac{Y}{Y+Z},$$

where $Y$ and $Z$ are the concentrations of the two species. Assuming independence of $Y$ and $Z$, the variance of $p_Y$ follows first-order error propagation:

$$\sigma_{p_Y}^2 = \left(\frac{\partial p_Y}{\partial Y}\right)^2 \sigma_Y^2 + \left(\frac{\partial p_Y}{\partial Z}\right)^2 \sigma_Z^2,$$

with partial derivatives

$$\frac{\partial p_Y}{\partial Y} = \frac{Z}{(Y+Z)^2}, \qquad \frac{\partial p_Y}{\partial Z} = -\frac{Y}{(Y+Z)^2}.$$

Therefore, the propagated standard deviation is

$$\sigma_{p_Y} = \sqrt{\frac{Z^2\,\sigma_Y^2 + Y^2\,\sigma_Z^2}{(Y+Z)^4}}.$$

The same approach extends to communities with more than two species by applying the multivariate delta method to $p_k = Y_k / \sum_\ell Y_\ell$ under an independence assumption.

##### S1.4 Uncertainty of Interaction Parameters

We detail the error calculation for growth rate interactions; the same procedure applies to final density interactions.

Define

$$a = \frac{X}{Y} - 1,$$

where $X$ is the growth rate measured in conditioned (spent) medium and $Y$ is the growth rate in fresh medium. Assume $X$ and $Y$ are independent random variables with means $\mu_x = \mathbb{E}[X]$, $\mu_y = \mathbb{E}[Y]$, and variances $\sigma_x^2 = \text{Var}(X)$, $\sigma_y^2 = \text{Var}(Y)$.

The expected value of $a$ can be expressed as

$$\mathbb{E}[a] = \mathbb{E}\left[\frac{X}{Y}\right] - 1 = \mathbb{E}[X]\,\mathbb{E}\left[\frac{1}{Y}\right] - 1,$$

and, under small variance in $Y$, approximated by

$$\mathbb{E}[a] \approx \frac{\mu_x}{\mu_y} - 1.$$

By the delta method, the variance of $a$ is approximated as

$$\text{Var}(a) \approx \left(\frac{\partial a}{\partial x}\right)^2 \sigma_x^2 + \left(\frac{\partial a}{\partial y}\right)^2 \sigma_y^2,$$

where the gradients evaluated at $(\mu_x, \mu_y)$ are

$$\frac{\partial a}{\partial x} = \frac{1}{\mu_y}, \qquad \frac{\partial a}{\partial y} = -\frac{\mu_x}{\mu_y^2}.$$

Hence,

$$\text{Var}(a) \;\approx\; \frac{\sigma_x^2}{\mu_y^2} + \frac{\mu_x^2 \sigma_y^2}{\mu_y^4},$$

and the corresponding standard deviation is

$$\sigma_a \;\approx\; \sqrt{\left(\frac{\sigma_x}{\mu_y}\right)^2 + \left(\frac{\mu_x \sigma_y}{\mu_y^2}\right)^2}.$$

#### S2 Theoretical Model of Microbiota Interactions

We model ecological interactions in well-mixed, liquid batch communities using a logistic formulation without spatial structure. The model was adapted from De Vos et al., 2017, with minor modification especially capping the final normalized population density on the double of its monoculture in fresh media.

Growth rate and final population density interactions affect the dynamic independently, such as

$$\frac{dX_i}{dt} = X_i\, g_i\, \text{max}\left(10^{-3},\, 1 + \sum_j a_{ij} X_j\right) \left(1 - \frac{X_i}{\text{min}\Big(\text{max}\Big(10^{-20},\, 1 + \sum_j b_{ij} X_j\Big)\,,\, 2\Big)}\right). \tag{2}$$

Here, $X_i$ denotes the abundance of species $i$, non-dimensionalized so that $X_i = 1$ equals the monoculture final density under our experimental conditions. The parameter $g_i$ is the intrinsic monoculture growth rate fitted directly from experimental growth curves.

The matrix $A = (a_{ij})$ quantifies interaction-mediated changes in the instantaneous growth rate of species $i$ due to species $j$. The growth multiplier is $1+\sum_j a_{ij} X_j$, so that $a_{ij} = 0$ indicates no detectable effect and, for $X_j \approx 1$, $a_{ij} = 1$ approximately doubles $i$'s instantaneous growth relative to $g_i$. The lower bound max$(10^{-3}, \cdot)$ prevents negative or zero multipliers under strong inhibition, ensuring numerically stable dynamics. Importantly, because equilibria satisfy the saturation term $(1 - X_i/K_i^{\text{eff}}) = 0$, growth-rate interactions primarily affect transient trajectories and establishment times, not the steady-state abundances.

The matrix $B = (b_{ij})$ modulates the effective carrying capacity of species $i$ in the presence of species $j$,

$$K_i^{\text{eff}} = \text{min}\left(\text{max}\left(10^{-20},\, 1 + \sum_j b_{ij} X_j\right),\, 2\right).$$

Thus $b_{ij} = 0$ implies no effect on the final density of $i$; for $X_j \approx 1$, $b_{ij} = 1$ doubles it, whereas sufficiently negative $b_{ij}$ compress $K_i^{\text{eff}}$ toward zero (competitive exclusion). The max bound enforces positivity and avoids numerical instability when interactions are strongly inhibitory; the outer min caps yields at twice the monoculture value, consistent with the normalization of $X_i$ and the empirical range of observed yields.

When $B \equiv 0$ (no carrying-capacity interactions) and interactions are weak, $K_i^{\text{eff}} = 1$ and the system converges to the monoculture yield $X_i \to 1$ as $t \to \infty$. Strongly negative entries of $B$ reduce $K_i^{\text{eff}}$ and can drive $X_i \to 0$, corresponding to exclusion or extinction.

##### S2.1 Comparison with an ecological variant

For reference, we also considered a variant closer to classical ecological models (Case, 1990, Hu et al., 2025), in which the carrying capacity enters linearly:

$$\frac{dX_i}{dt} = X_i\, g_i\, \mathrm{max}\left(10^{-3},\, 1+\sum_j a_{ij}X_j\right)\left(\mathrm{min}\left(1+\sum_j b_{ij}X_j,\, 2\right) - X_i\right). \tag{3}$$

In the main model (Eq. 2), the saturation term is dimensionless, $1 - X_i/K_i^{\text{eff}}$, with $K_i^{\text{eff}}$ normalized to the monoculture yield. The ecological variant (Eq. 3) uses the traditional $X_i(K_i - X_i)$ form with $K_i^{\text{eco}} = \mathrm{min}(1 + \sum_j b_{ij}X_j, 2)$. Both formulations behave similarly when interactions are weak and $K_i \approx 1$; however, under strong inhibition the normalized form yields smoother transitions and avoids abrupt changes associated with near-zero $1 + \sum_j b_{ij}X_j$.

##### S2.2 Computation of community endpoints

We determined long-term community outcomes by directly computing fixed points of the steady-state equations. For efficiency during sensitivity analyses, steady states were obtained by setting the right-hand side of Eq. (2) to zero and solving the resulting algebraic system with `scipy.optimize.fsolve`, constrained to solutions consistent with the carrying-capacity term.

A community was classified as stable if every species present in the initial inoculum reached a final abundance greater than or equal to its inoculum level. This criterion reflects the empirical observation that species remaining at or below their inoculum concentration typically fail to persist upon passage, whereas those exceeding it reliably establish.

For comparison with a model without interaction parameters (Supplementary Fig. 8), we also performed numerical time integration of Eq. (2).
The system was integrated for $t = 24$ h using `scipy.integrate.solve_ivp` with the stiff `BDF` method. The ODE system included only the species present in the initial inoculum vector. The final state $X_i(t_{\text{end}})$ was taken as the predicted end point and the communities were classified as stable using the same criterion described above.

#### S3 Invasion Assay Using the Theoretical Model

##### S3.1 Simulating invasion into previously stabilized communities

We quantified the effect of introducing a novel microbial species into resident communities that had converged to a stable state in the baseline analysis. Simulations were implemented in Python and parallelized using `joblib.Parallel`.

For each resident community, we recovered the species composition and the corresponding final abundances. We then expanded the community by adding a single invader with a small initial inoculum set to 0.05 of its final density of monoculture (i.e. $x_{\text{inv}}(0) = 0.05\,\hat{X}_{\text{inv}}$). Resident species were initialized at their previously computed final abundances. Model parameters for resident species were taken from the same sensitivity-analysis draw used to identify that resident community. Invader parameters were sampled as described in the next subsection.

The augmented system was integrated up to $t = 48$ h using a stiff ODE solver (`solve_ivp` in Python, method `BDF`). The full community dynamics followed the same theoretical ecological model used in the stability analysis, but now including both the resident species and the introduced invader. The integration time of $48\,$h was chosen to ensure that both the establishment of the invader and the potential responses of resident species to be reliably captured.

To interpret the simulation outcomes, we compared each species' final abundance to its corresponding initial inoculum. For the invader, the threshold was its inoculum in the augmentation experiment. For resident species, we compared the final abundances to the inoculum used in the original stability experiments, rather than to the abundances of the previously identified stable communities.

Based on these comparisons, invasion outcomes were classified into four categories:

- **Augmentation:** The invader successfully establishes (final abundance exceeds its inoculum), and all resident species remain above the stability threshold.
- **Displacement:** The invader establishes, but at least one resident species falls below the stability threshold.
- **Resistance:** The invader fails to establish (final abundance remains at or below inoculum), while all resident species remain above the stability threshold.
- **Disruption:** The invader does not establish, and at least one resident species falls below the stability threshold, indicating destabilisation of the resident community.

##### S3.2 Generation of random invader species

To assess how resident communities and invader characteristics modulate invasion outcomes, we generated 500 synthetic invader species. For each invader, all model parameters were sampled independently from Normal distributions fit to experimental measurements.

- **Monoculture traits (fresh medium):** The intrinsic growth rate $g_{\text{inv}}$ and the final population density $\hat{X}_{\text{inv}}$ were sampled from normal distributions fitted to the observed monoculture growth and final density across the experimental results.
- **Interaction coefficients:** Growth-rate interactions ($A = (a_{ij})$) and final-density interactions ($B = (b_{ij})$) were sampled from normal distributions fitted to the experimentally measured interaction strengths among the selected bacterial species. Because each parameter is sampled independently to avoid introducing correlations into the model, we generated all invader–resident interaction parameters from fitted normal distributions derived from the experimental data:
  - **Resident-to-invader effects:** for each resident $j$, we sampled $a_{\text{inv},j}$ and $b_{\text{inv},j}$, quantifying the invader's response to medium conditioned by resident $j$.
  - **Invader-to-resident effects:** for each resident $i$, we sampled $a_{i,\text{inv}}$ and $b_{i,\text{inv}}$, quantifying each resident's response to medium conditioned by the invader.

The parameters of the invaders were independently sampled across traits and species to avoid adding correlations that could make the invasion frequencies harder to interpret. Some empirical distributions showed bimodality, but we used normal distributions based on the observed mean and standard deviation. This approach gives a smooth, conservative approximation that avoids overfitting to the limited set of bacterial strains while still matching the main features of the data.
